# Genetic sex determination in three closely related hydrothermal vent gastropods, including one species with intersex individuals

**DOI:** 10.1101/2023.04.11.536409

**Authors:** J Castel, F Pradillon, V Cueff, G Leger, C Daguin-Thiébaut, S Ruault, J Mary, S Hourdez, D Jollivet, T Broquet

## Abstract

Molluscs have a wide variety of sexual systems and have undergone many transitions from separate sexes to hermaphroditism or vice versa, which is of interest for studying the evolution of sex determination and differentiation. Following the serendipitous observation that sex was the primary driver of genetic structure in the hydrothermal vent gastropod *Alviniconcha boucheti*, we investigated sexual systems and sex determination in this species and two others of the same genus. We combined genome-wide multi-locus genotypes obtained from RAD sequencing with anatomical observations of the gonads of the three *Alviniconcha* species occurring in the southwest Pacific Ocean: *A. boucheti* (*n*=199), *A. strummeri* (*n*=41 ind.) and *A. kojimai* (*n*=246). In two of the species (*A. boucheti* and *A. strummeri*), the sexes are separate and genetically determined by a male-heterogametic (XY) system. External observation of the gonads in the third species (*A. kojimai*) also suggested that the sexes were separate, but histological analyses revealed that 76% of the individuals classified as females from external observation of the gonads presented a mosaic of male and female reproductive tissue. Empirical analyses and simulations showed nonetheless that 14 RAD loci were sex-linked with an XY signature in *A. kojimai* (as compared with 64 in *A. strummeri* and 373 in *A. boucheti*). Comparison across species and mapping of RAD loci to a non-contiguous reference genome assembly of the related species *A. marisindica* showed that all sex-linked loci identified in *A. kojimai* are located on five scaffolds that also contain 15 and 67 sex-linked RAD loci in the other two species, respectively. These results suggest that all three *Alviniconcha* species share the same XY sex determination system, but that the gonad of XX *A. kojimai* individuals are invaded by a variable proportion of male reproductive tissue. It remains to be seen whether the male tissue in these intersex individuals is functional or not. The identification of Y-specific RAD loci (found only in *A. boucheti*) and the phylogenetic analysis of three sex-linked loci shared by all species suggested that X-Y recombination has evolved differently within each species. This situation of three species showing variation in gonadal development around a probably common sex determination system provides new insights into the reproductive mode of poorly known deep-sea species and questions the evolution of gametogenetic polymorphism in these species.

## Introduction

A wide variety of sexual systems are observed in eukaryotes, raising questions about the evolution of this polymorphism (recent reviews in Leonard, 2018; Pannell & Jordan, 2022). For instance, male and female gametes are produced by different individuals in about 95% of animal species (gonochoric species, Jarne & Auld, 2006), while an almost exact opposite figure was estimated for angiosperms (dioecious species, Renner, 2014). On closer examination, the seemingly simple, contrasted situation of animals and plants hides a diversity of sexual systems scattered within each of these broad taxonomic groups (and this diversity of course extends to other regions of the tree of life, e.g. Coelho & Umen, 2021). Considering animals, hermaphroditism (defined as a sexual system where male and female reproductive organs are simultaneously or sequentially found on the same individual) is nearly absent in insects but widespread across many other groups where it occurs in variable frequency (Jarne & Auld, 2006; Leonard, 2018). This means that hermaphroditism and gonochorism have evolved multiple times, in a wide variety of phylogenetic lineages. Some large clades (e.g. insects, tetrapods, chondrychtyes) have a mostly invariant, stable sexual system (Leonard, 2018; Pannell & Jordan, 2022), but others show much polymorphism. For instance, Jarne and Auld (2006) estimated that hermaphroditism evolved from gonochorism at least 40 times within prosobranch gastropods alone, raising questions as to how and why such transitions occur.

Evolutionary transitions between sexual systems are intimately linked with the evolution of sex determination and sex differentiation. Sex determination and differentiation can vary from purely genetic, when variation at some position in the genome triggers the male or female specific developmental program, to purely epigenetic, when sex-specific programs are activated by differential expression (Beukeboom & Perrin, 2014). A variety of determination systems along this continuum may lead to separate sexes (gonochory), but hermaphroditism is normally found towards the epigenetic end of the continuum (Beukeboom & Perrin, 2014). Although the evolution of sex determination is a lively field of research, many questions remain regarding the evolution of sex determination mechanisms involved in transitions between gonochory and hermaphroditism and the directionality of these changes in animals (Leonard, 2018), and also in the cost and evolution of complex sexual systems involving for instance a mixture of hermaphroditic individuals and males (androdioecy) or females (gynodioecy). Separate sexes may evolve *de novo* from hermaphroditism through a series of mutations that create one sex and then the other one, followed by selection for linkage between these loci (that is, formation and evolution of sex-chromosomes). This process and its molecular bases have been described in angiosperms (reviewed in Charlesworth, 2006) and have recently been investigated in Platyhelminthes and nematodes for instance (Wang *et al*., 2022). In the reverse direction, transitions from separate sexes to hermaphroditism can also be achieved through a series of mutations affecting the sex determination pathway that will allow individuals of one sex to become hermaphrodites, which may provide them with a fitness advantage when access to mating partners is limited. For instance, self-fertile hermaphrodites have evolved from XX females following different modifications of the core sex determination cascade in *Caenorhabditis* species (therefore resulting in androdioecious systems, e.g. Kiontke *et al*., 2004; Hill *et al*., 2006).

Here we report the serendipitous discovery of genetic sex determination in three species of deep-sea hydrothermal vent symbiotic gastropods (genus *Alviniconcha*), as well as the unanticipated observation that one of these species contains individuals with a mixture of male and female reproductive tissue, therefore providing an interesting case of a mixed or transitory sexual system.

Gastropods exhibit a wide range of sexual systems and sex determination systems whose evolution has involved a high frequency of transitions (Jarne & Auld, 2006; Collin, 2013; Leonard, 2018). To our knowledge there is no known case of functional andro-or gynodioecy in this group (Weeks, 2012) but gastropods have been particularly studied in the context of exogeneous, anthropogenic, effects on intersex (defined in molluscs as the simultaneous presence of oocytes or spermatogonia, at varying degrees of development, within the normal gonad of the opposite gender of a gonochoristic species, Grilo & Rosa, 2017). Sex determination and sex differentiation of gastropods as a group seem to be very labile, which is interesting for understanding the evolution of these systems. Although marine gastropods are primarily gonochoristic (reviewed in Collin, 2013), this is not a universal rule (e.g. simultaneous hermaphroditic Ophistobranchia, sex-changing Patellidae and Calyptreidae). In particular, we know very little about the biology of many deep-sea species because these environments are difficult to access. Hydrothermal vents are found along tectonically active locations such as mid-oceanic ridges and active volcanic arcs where seawater penetrates through faults in the lithosphere and resurfaces as a hot, anoxic, acidic fluid enriched in metals and dissolved gases. The chemical and physical characteristics of these peculiar environments have led to specific adaptations, including the establishment of symbioses with chemoautotrophic bacteria (Cavanaugh, 1983; Childress, 1995), and constraints on immune systems (Papot *et al*., 2017), respiratory systems (Hourdez & Lallier, 2007), detoxification systems (Powell & Somero, 1986), and dispersal allowing the colonization of fragmented and ephemeral habitats (Lutz, 1988; Vrijenhoek, 2010). Regarding reproductive strategies, the hydrothermal environment seems to have generally favoured continuous gametogenesis, oocyte and sperm storage, internal fertilisation, and lecithotrophic development based on yolk reserves (Tyler & Young, 1999; Jollivet *et al*., 2000; Faure *et al*., 2007), although some abundant species do not follow this broad pattern (e.g. high fecundity and planktotrophic larvae in *Bathymodiolus* mussels, Laming *et al*., 2018, and *Alviniconcha* gastropods). The sexual system of many hydrothermal vent species, not to mention their sex determination system, remains to be elucidated.

Here we studied *Alviniconcha* gastropods, large hairy snails that form dense aggregations in warm (7-42°C), sulphide-rich (up to 250 µmol.l^-1^), and low-oxygen (< 50 µmol.l^-1^) habitats associated with hydrothermal vents (Desbruyères *et al*., 1994; Podowski *et al*., 2010). There are six species in the genus (Johnson *et al*., 2015; Breusing *et al*., 2020; Castel *et al*., 2022), all forming symbiotic associations with sulfo-oxydizing chemoautotrophic bacteria in their gills (Stein *et al*., 1988). We focus here on *Alviniconcha boucheti*, *A. strummeri*, and *A. kojimai*, three species that inhabit the hydrothermal vents of the back-arc basins in the Western Pacific Ocean.

Following the unexpected observations of i) sex-linked patterns of genetic structure and ii) a proportion of male and female tissue mixed within the gonad of some individuals in *A. kojimai*, the aims of this study were, firstly, to identify the sexual system and sex determination in the three species and, secondly, to assess whether the genomic position of sex-linked loci is conserved across species to better understand the apparent flexibility of sexual systems in this genus.

## Methods

### Sampling and sexing

We used a subset of a previously published genomic data set of ddRAD-seq sequences obtained by Castel et al. (2022). In short, here we used data from 199 *A. boucheti*, 246 *A. kojimai*, and 41 *A. strummeri* individuals (total 486) sampled from 10 vent fields in four back-arc basins and a volcanic arc of the Western Pacific Ocean (Figure 1; Table S1). All samples were collected during the Chubacarc expedition on board of the R/V L’Atalante with the ROV Victor 6000 in Spring 2019 (Hourdez & Jollivet, 2019). Details about sampling conditions and the preparation of samples on board are described in Castel *et al*. (2022).

**Figure 1:**
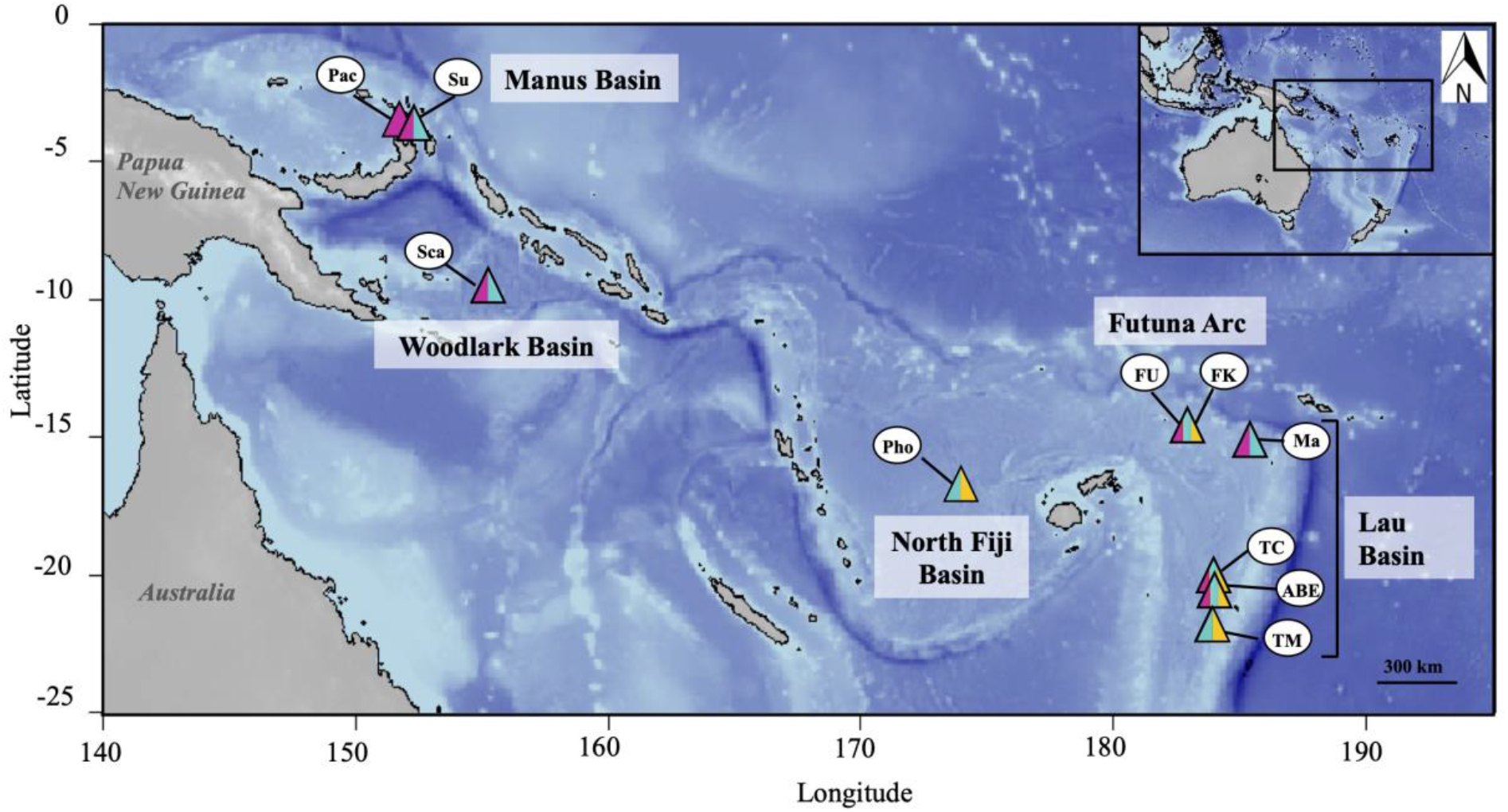
Origin of samples. Manus Basin: Susu (Su) and Pacmanus (Pac). Woodlark Basin: La Scala (Sca). North-Fiji Basin: Phoenix (Pho). Futuna volcanic arc: Fatu-Kapa (FK) and Fati-Ufu (FU), these two sites are indistinguishable at this scale. Lau Basin: Mangatolo (Ma), Tow Cam (TC), ABE (ABE), and Tu’i Malila (TM). Species *A. boucheti* is represented in purple, *A. kojimai* in turquoise, and *A. strummeri* in yellow.

As no sexual dimorphism is externally visible in *Alviniconcha* gastropods, we determined the sex of individuals from observations of the gonads using a two-step approach. First, sex was identified from the external colour of gonads for a fraction of individuals that were dissected on-board during the sampling cruise (268 out of the 486 ind. that have been processed, 55%). The presence of oocytes gives a whitish colour to the gonads that indicates female sex while brown gonadal tissue is specific to males, but this method only allowed a rough identification of the sex, with some uncertainty. Second, a sub-sample of 166 individuals (34%) were studied using histological observations of the gonad (including 163 individuals previously sexed on board from gonad colour). A cross section of the central part of the formalin-fixed gonad was taken and embedded in paraffin. Sections 5-µm thick were then cut using a Leica RM2035 microtome and stained with standard histological protocols for Harry’s haematoxylin and eosin. The stained sections were then observed using a Leica DM2500 LED binocular microscope. The sex-ratio was not estimated from these internal observations of the gonads because these individuals were not strictly randomly chosen onboard (we tended to balance the number of males and females selected for dissection to study the gametogenesis in the three species).

### Genotyping

We used the raw sequence data obtained in (Castel *et al*., 2022). As detailed in this previous article, DNA was extracted from 570 foot tissue samples using NucleoSpin® Tissue kits (Macherey-Nagel) or a CTAB protocol (Doyle & Dickson, 1987; modified in Jolly *et al*., 2003), and individual-based ddRAD libraries were produced following the protocol detailed in Daguin-Thiébaut et al. (2021). The libraries were sequenced in 150-bp paired-end reads using a Novaseq 6000 (library construction and sequencing detailed in Castel *et al*., 2022). Raw reads were demultiplexed and paired using the process_radtags module of Stacks software version 2.52 (Catchen *et al*., 2011, 2013).

These demultiplexed reads were then re-analysed for the present study. We used Kraken v.2 (Wood & Salzberg, 2014) to remove potential reads originating from chemoautotrophic bacteria and keep only the eukaryotic reads for the assembly (between 0.9% and 15% of the initial reads assigned to prokaryotes were discarded, depending on the species considered). The assembly parameters were chosen independently for each species (unlike in Castel *et al*., 2022) on the basis of pilot studies consisting of a systematic exploration of the effect of the main Stacks parameters *m*, *M*, and *n.* Details of the pilot analyses and final conditions of data treatment are presented in the supplementary material.

The SNPs identified by Stacks were filtered in R version 4.1.1 (R Core Team, 2021) to keep the SNPs present in at least 90% of the individuals and only keep the individuals which were genotyped at least at 80% of the total number of SNPs (allowing a maximum of missing data of 20% for a given individual), except in *A. strummeri* for which we had less samples (because this species has a smaller range and lower abundance) and thus permitted more missing data (20% per SNP and 40% per individual). A total of 499 individuals passed these filters (163 117 SNPs in *A. boucheti*, 70 122 SNPs in *A. kojimai*, and 36 568 SNPs in *A. strummeri*). The distribution of the mean depth of coverage per SNP in these datasets is detailed in supplementary material (figures S4 and S5).

We visualized the genetic structure of each species using principal components analyses in R (adegenet package, Jombart, 2008) based on a single SNP per RAD locus (table 1). This analysis also allowed us to further filter out eight samples in *A. boucheti* and five in *A. kojimai* that had a strong effect on the PCA (pairs of individuals driving in turn the PCA, potentially due to high relatedness, or contamination). Our final data set therefore contained 199 *A. boucheti*, 246 *A. kojimai,* and 41 *A. strummeri* (total 486 ind., table S1).

**Table 1.** Diversity indices for the three species of *Alviniconcha. H*_T_: total gene diversity, *H*_S_: sex-specific gene diversity, where *m* is for male and *f* stands for female in *A. boucheti* and *A. strummeri*, and either female or morphological hermaphrodites in *A. kojimai* (see main text); *F*_ST_: differentiation between sexes; *H*_O,*i*_: observed heterozygosity in sex *i* and *F*_IS,*i*_: heterozygosity deficiency in sex *i*. Column “Unsexed” gives the number of individuals with uncertain or unknown phenotypic sex (none of these individuals were used in diversity calculations). These statistics were computed using the samples that passed all filtering steps.

| Species | Males | Females | Unsexed | Total | SNPs | $H_T$ | $H_{S,m}$ | $H_{S,f}$ | $F_{ST}$ | $H_{O,m}$ | $H_{O,f}$ | $F_{IS,m}$ | $F_{IS,f}$ |
| --- | --- | --- | --- | --- | --- | --- | --- | --- | --- | --- | --- | --- | --- |
| *A. boucheti* | 51<sup>a</sup> | 46<sup>a</sup> | 102<sup>a</sup> | 199 | 96 768 | 0.127 | 0.127 | 0.126 | 0.01 | 0.127 | 0.124 | 0.001 | 0.013 |
| *A. strummeri* | 10 | 12 | 19 | 41 | 25 996 | 0.257 | 0.258 | 0.256 | 0.002 | 0.251 | 0.247 | 0.028 | 0.034 |
| *A. kojimai* | 81 | 68 | 97 | 246 | 33 576 | 0.098 | 0.098 | 0.098 | 0.0008 | 0.098 | 0.097 | 0.008 | 0.011 |

<sup>a</sup> These numbers take into account the fact that three *A. boucheti* individuals with inconsistent phenotypic vs genotypic sexes were reclassified as "Unsexed" (see text).

Because sex appeared to be a key driver of genetic structure (see Results), the distribution of genetic diversity was further investigated using functions in the R package hierfstat (Goudet & Jombart, 2021), using only the sexed individuals (97 *A. boucheti*, 149 *A. kojimai*, and 22 *A. strummeri*, table 1), and again, a single SNP per RAD locus. We characterised the global genetic diversity within each species by estimating total gene diversity *H*_T_. The diversity within each sex *i* was then described by calculating the gene diversity *H*_S,*i*_ and observed heterozygosity *H*_O,*i*_, and the discrepancy between these parameters (indicating a potential departure from random association of gametes) was estimated as *F*_IS_ within each sex. Finally, the genetic differentiation between sexes was estimated using Weir and Cockerham (1984) estimator of *F*_ST_ between males and females and tested for statistical significance using 100 bootstraps over loci in StAMPP (Pembleton *et al*., 2013).

### Identification of the genetic sex determination system

Identifying sex chromosomes can be achieved by looking for loci that are present on only one of the sex chromosomes (Gautier, 2014; Stovall *et al*., 2018; Feron *et al*., 2021; Garcia-Raventós *et al*., 2023), or by looking at heterozygosity and differentiation between sexes at homologous loci (e.g. Berset-Brändli *et al*., 2006; Brelsford *et al*., 2017; Käfer *et al*., 2021; Hearn *et al*., 2022; see also Pyne *et al*., 2022 for an association study). Here we followed the second approach (see Palmer *et al*., 2019 for a review of methods). To detect genetic systems of sex determination, we searched for SNP markers with alleles differentially fixed on sex chromosomes. If sex was genetically determined and if some SNPs in a non-recombining region had alleles that were differentially fixed on sex chromosomes, then these SNPs would be strictly homozygous in the homogametic sex (XX females or ZZ males) and heterozygous in the heterogametic sex (XY males or ZW females). We looked for such SNPs by estimating the observed heterozygosity in males *H*_O,m_ and females *H*_O,f_ at each SNP using functions from the hierfstat R package (Goudet & Jombart, 2021).

### Identification of sex-linked loci

Besides sex-linked SNPs with fixed, divergent alleles on sex chromosomes, other polymorphic SNPs located on sex chromosomes may have various levels of heterozygosity and sex-linked differentiation depending on the genomic landscape of recombination along these chromosomes (see, for example, Muyle *et al*., 2017 for a related discussion). Hereafter, ‘sex-linked’ will refer to any pattern of genetic variation that is not independent of sex, including polymorphism segregating on regions of the sex-chromosomes undergoing partial recombination or recent recombination arrest. The distribution of genetic diversity between sexes (*F*_ST_) and between individuals within each sex (*F*_IS_) is more informative than *H*_O_ to detect sex-linked loci that are not differentially fixed, because sex linkage will tend to result in an excess of heterozygotes in the heterogametic sex (e.g. XY males) and a simultaneous increased differentiation between sexes.

However, we expect some variance associated with these statistics measured using a finite number of male and female individuals (Table 1). Therefore, for each species we computed *F*_IS_ and *F*_ST_ values from simulated genotypes randomly created from a range of allelic frequencies assuming autosomal segregation for two groups of individuals with sample sizes equal to the actual size of the male and female groups in our real dataset (thereby introducing estimation of error variance in the simulations). The simulations were coded in R and consisted of the following steps:

- Create a number of theoretical bi-allelic SNPs equal to the number of SNPs genotyped in each species and where each SNP has a reference allele with frequency *p* and an alternative allele with frequency 1-*p*, where *p* was randomly sampled from a uniform distribution between 0 and 1 (such as to cover all possible initial values for allelic frequencies).
- Create a group of “male” and a group of “female” genotypes (with sample sizes equal to the number of males and females sampled in each species) by sampling two alleles randomly from a binomial distribution parameterized using the expected allelic frequencies simulated in step 1.
- From these simulated genotypes, estimate *F*_ST_ between the “male” and “female” groups, and *F*_IS_ within each of these groups at each simulated SNP. These estimates were calculated following (Nei, 1987) as described in R package hierfstat and simplified for bi-allelic SNPs:

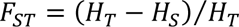

and

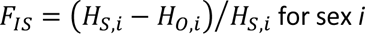

where:

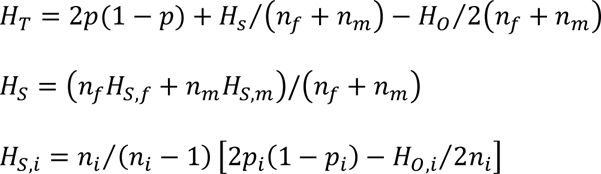

with *p* the allelic frequency of the reference allele (recalculated from the simulated genotypes) at the SNP under consideration, *n*_*i*_ is the sample size of sex *i*. In these equations, 2*p*(1-*p*) is the gene diversity expected under random mating (Hardy-Weinberg, autosomal segregation at a bi-allelic loci with allelic frequencies *p* and (1-*p*)), and all other terms are bias corrections for small sample sizes as described in Nei (1987).

These statistics were computed for each SNP. After simulation, observed *F*_ST_ and sex-specific *F*_IS_ values at each SNP from our dataset can be compared to the simulated values. All SNPs from the real dataset that had combinations of *F*_IS_ and *F*_ST_ values outside of a convex envelope encompassing all possible theoretical values obtained from autosomal simulations had properties that were not compatible with autosomal segregation. The convex envelope was built using slightly modified R code from F. Briatte (https://github.com/briatte/moralecon/blob/master/moralecon.r).

### Identification of loci present only on the Y chromosome

The filters for missing data described above were designed to obtain high-quality SNP genotypes. However, since we found sex determination to be encoded by a XY system (see results), our filters may have discarded RAD sequences contained in genomic regions that would be present only on the Y chromosome (as such loci would be missing from all XX individuals, i.e., in our case half of the individuals in each species). Hence, we ran another set of analyses using sex-specific filters for missing data so as to search for potential Y-specific RAD loci. Specifically, we looked for SNPs totally absent from females while being genotyped in at least 10 males, therefore pointing to genomic regions that are specific to the Y, or homologous but strongly divergent on X and Y chromosomes (and therefore identified as two different loci by the Stacks pipeline). These analyses were repeated for each species.

### Synteny of sex-linked loci across the three species

As we used different SNP catalogues for each species in Stacks, sex-linked RAD loci identified within each species were compared to each other through reciprocal blastn (e-value threshold= 10^-5^, word size=28) in Geneious (Kearse *et al*., 2012) in order to identify orthologous RAD loci between two or three species.

In a second step, we search for the position of all sex-linked RAD sequences in the closest reference genome available (*A. marisindica*, assembly ASM1885773v1, Yang *et al*., 2022). Although the level of divergence expected between *A. marisindica* and the species studied here prevented us from using this genome for processing RAD-sequences (we used a de-novo approach instead, see methods in Castel *et al*., 2022) and although this reference genome is not contiguous (3926 scaffolds, N50 = 0.73 Mb, L50 = 336 contigs), it is still useful to identify potential RAD sequences that would be obtained in only one or two of the three species but belonging to a single genomic scaffold. To this end, we ran a blastn analysis using the software Geneious to identify the best hit of each sex-linked RAD locus on the *A. marisindica* assembly, with minimum coverage of 95% of the RAD sequence and an e-value threshold of 10^-5^. We chose these conservative parameters in order to obtain reliable positive hits and to detect scaffolds that are undoubtedly sex-linked, at the cost of missing additional scaffolds that could contain sex-linked RAD loci (loci contained in repeated elements that occur on multiple scaffolds or more diverging sequences with less than 95% coverage). We therefore aimed to obtain an estimate of the minimum size of sex-linked regions in each species.

### Phylogeny of sex-linked RAD haplotypes

To investigate the evolutionary history of *Alviniconcha* sex chromosomes, we focused on the three RAD loci that were found to be sex-linked in all three species (see results and Fig. 6A; these loci hereafter designated locus 1-3 were 218 to 385 bp long). We used the individuals for which sex was identified from histological analyses, that is, 9 to 43 XY individuals (males) and 12 to 40 XX individuals (females or morphological hermaphrodites, see results) per species (Fig. 7). The pairs of sequences obtained for each diploid individual at a given RAD locus were aligned in MEGA v. 11.0.13 (Tamura *et al*., 2021), and then used to construct a phylogenetic UPGMA tree based on Felsenstein’s (1981) distances with R packages ape v5-7.1 (Paradis & Schliep, 2019) and phangorn 2.11.1 (Schliep, 2011). The trees obtained show the phylogeny of 290 to 310 sequences (bootstrap values based on 100 resampled datasets), each of which was either located on an X or Y chromosome. All sequences clustered into 6 to 11 haplotypes, and the association of each sequence to the X or Y chromosome was deduced from the frequency of haplotypes in each sex. This deduction was facilitated here by the fact that in nearly all cases the males of a given species shared only one haplotype that was absent from all females. This was not the case for the third locus in *A. kojimai* (Fig. 7 C), and so Inferring the X or Y location of the sequences was less straightforward. In this case, we first calculated the frequency of each haplotype on X chromosomes using XX individuals only. Then, we calculated the frequency of each haplotype on Y chromosomes using XY individuals only (knowing the frequency of each haplotype on X).

## Results

### Gender identification and sex ratio

The putative sex of *A. boucheti* individuals was determined on-board by external observation of the gonads for 98 individuals, and later reassessed in the lab by histological observations for 52 of these individuals and two others (i.e. 100 individuals examined in total). Of the 52 possible comparisons between the two observations, only 5 mismatches were observed. That is, sex identification based on external observation of the gonads generated about 9.5% error. Combining the two datasets (as reported in Table S1), we obtained 47 females and 53 males.

For *A. strummeri*, 22 individuals were sexed on-board, and all of them were also used for gonad cross-sections. We observed 2 discrepancies, that is, again 9% error from external gonad observation. Histological observations indicated that we had 12 females and 10 males.

In *A. kojimai*, 67 females and 81 males were identified on-board. Surprisingly, from the subset of 41 females later used in histological analyses, 31 presented a gonad cross-section that included some proportion of male tissue, as shown in figure 2. Visual inspection of gonad sections from these individuals revealed variable proportions of male tissue. By contrast, histological observations confirmed the sex of 47 out of the 48 males for which we took gonad cross-sections (the remaining individual was a female wrongly identified from external observation of the gonad, which brings the total number of females identified by histology to 11).

**Figure 2:**
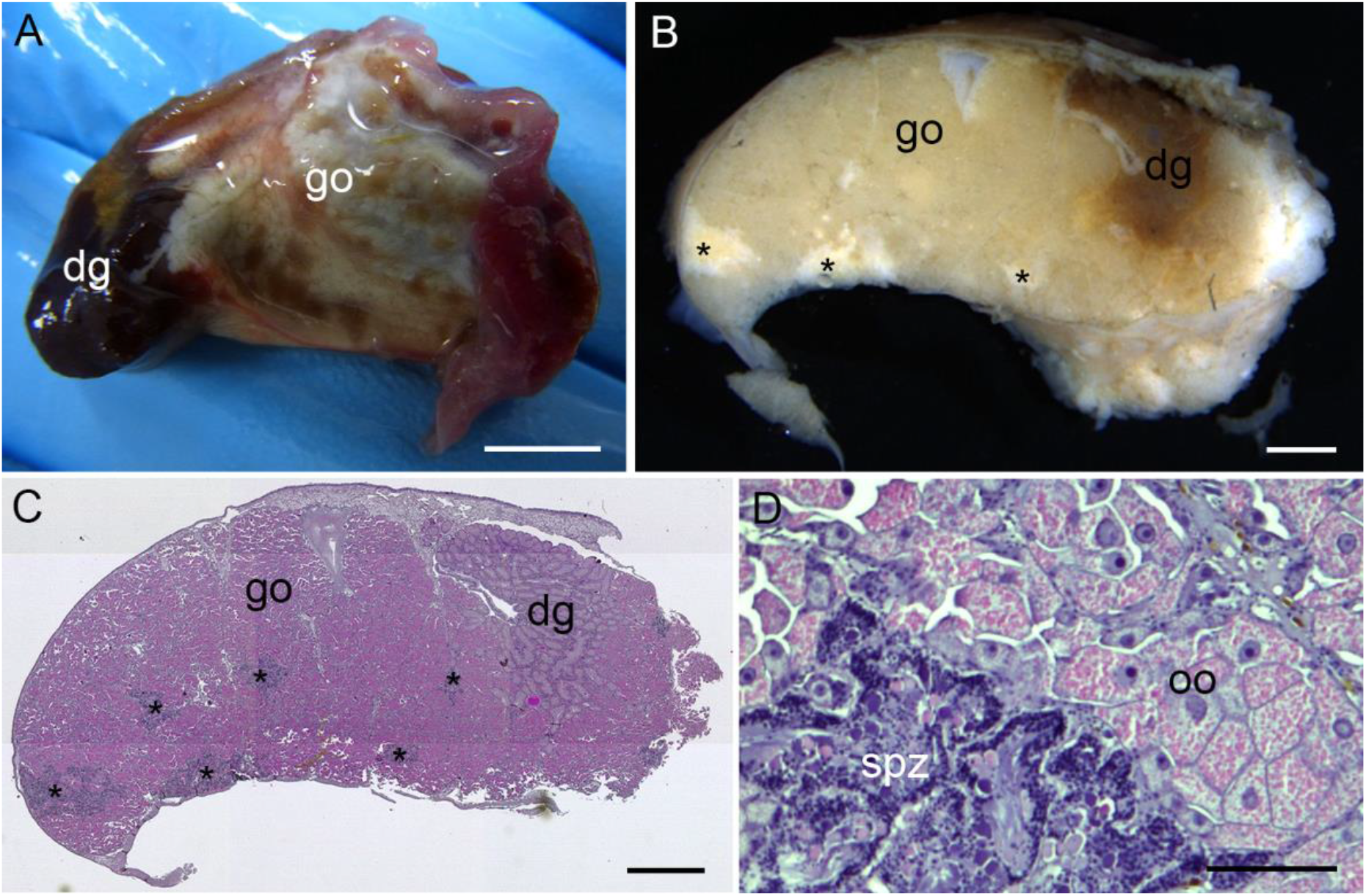
Gonad of an *A. kojimai* ‘morphological hermaphrodite’. A) Gonad removed from fresh tissue on board. Note its overall whitish colour used as a diagnostic for the female sex on board, while the brownish spots scattered over it reflect male tissues. B) Transverse cross-section of a formalin-fixed gonad (same individual as in A), taken approximately at the middle of the antero-posterior axis. Asterisks indicate male tissue. C) Hematoxylin-eosin coloured semi-thin section prepared from the section shown in B where male tissues appear in blue (also marked with asterisks) and female tissues appear in pink. D) Close-up of gonadal tissues with both male and female gametes developing in close proximity. dg: digestive gland, go: gonad, oo: oocytes, spz: spermatozoa. Scale bars are 5 mm in A, 1 mm in B & C, 100 µm in D.

Focusing on data from histology only, we identified 48 males, 11 females, and 31 ‘morphological hermaphrodites’ or ‘intersex individuals’ (which we define as individuals with coexisting male and female gonadal tissue but no information on the functionality of these tissues, see discussion).

Because our histological observations were limited to a single middle cross section per individual, it is possible that some male tissue was missed in the individuals identified as females (see discussion). Therefore, unless otherwise stated, we considered two groups of *A. kojimai* individuals in downstream analyses, based on their sex as identified from gonad sections when available (on-board observations when not available): a group of males (*n* = 81) and a group of ‘females or morphological hermaphrodites’ (*n* = 68). See the discussion for a detailed argumentation about this dichotomy.

### Distribution of genetic variation

The principal components analysis of SNP genotypes in *A. boucheti* revealed an unexpected pattern (Fig. 3A): the first axis (1% of the total variance) separated two groups of individuals clearly determined by their sex, except for three individuals that segregated with the opposite sex. Two of these three individuals had been processed consecutively on-board and their numbering was most likely inverted by mistake. The sex of the third one was identified from external observation only, and given the rate of error of this method it is possible that sex was wrongly identified. All other sexed individuals (i.e. 46 females and 51 males) clustered into two distinct groups along the axis 1 of the PCA. In addition, the 99 unsexed individuals were evenly distributed between the two clusters (50 with the male cluster and 49 with the female cluster, Fig. 3A). Using axis1 coordinates to identify the sex of all individuals, we find 101 males and 95 females (*r* = 0.51, exact binomial test *p*=0.72). The second axis (0.8% of the total variance) further subdivided the male and female groups according to the main geographical subdivision of sampling locations: Western sites (Manus and Woodlark basins) versus Eastern sites (Lau basin and Futuna arc), which are separated by well over 2000 km (Fig. 1).

**Figure 3:**
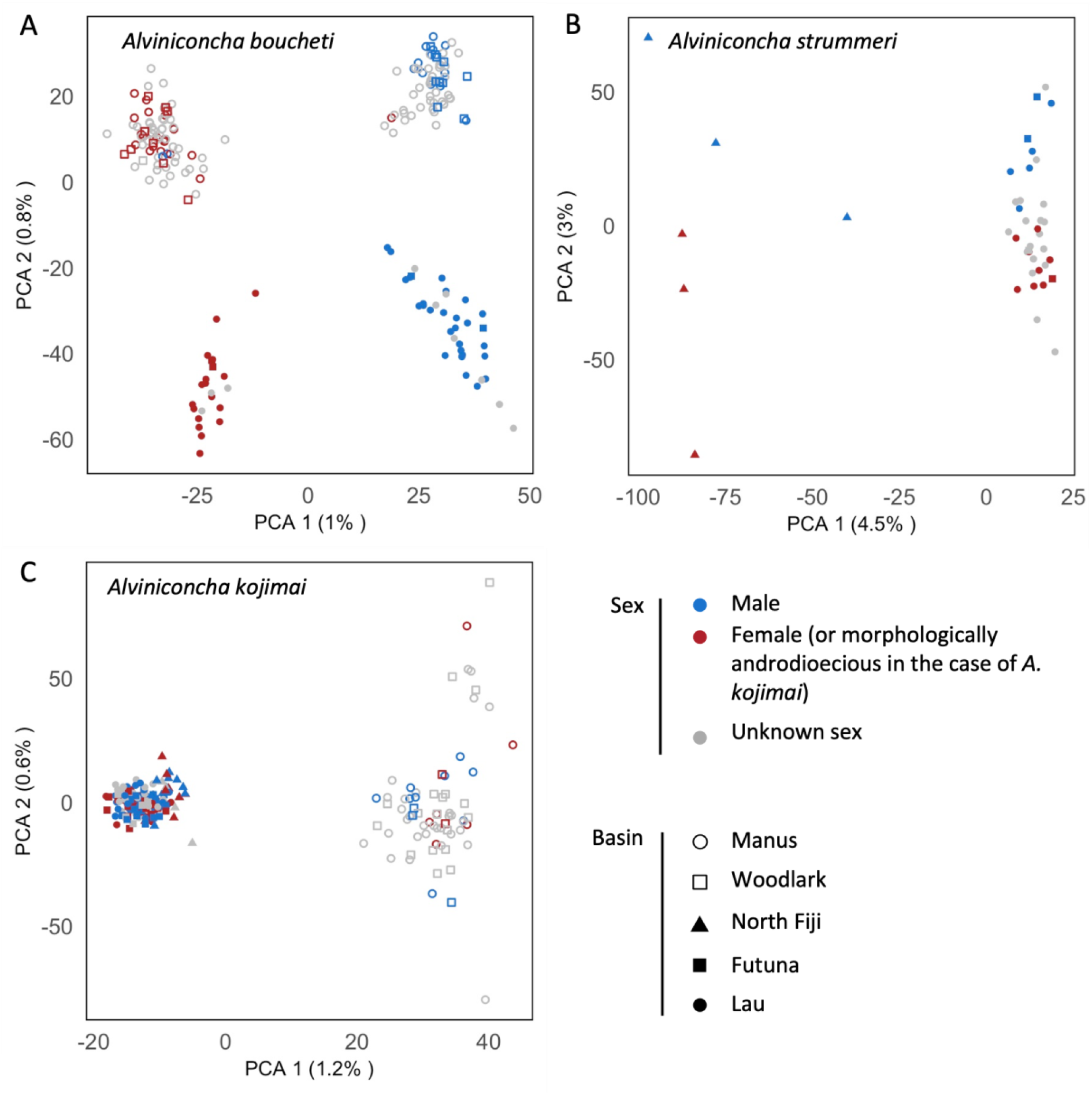
Principal components analyses of SNP genotypes for the three species of *Alviniconcha* gastropods. Solid symbols correspond to the Eastern basins, while empty symbols correspond to the Western basins (Figure 1). A) *A. boucheti* (199 individuals, 96 768 SNPs). The first axis separates individuals according to their sex, while the second axis is consistent with geography. B) *A. strummeri* (41 individuals, 25 996 SNPs). While the first axis separates individuals according to their geographic origin, the second axis is driven by sex. C) *A. kojimai* (246 individuals, 33 576 SNPs). The first axis separates individuals according to their geographic origin. There was no obvious clustering by sex in this species.

Remarkably, sex was also a key driver of genetic variation in *A. strummeri* (Fig. 3B), but with a more limited effect since it separated individuals along the second component of the PCA (3% of the total variance), while the geographical distribution of sites explained the distribution of genotypes along the first axis (4.5% of the total variance, despite the smaller geographic range of this species). Although less clear-cut than in the case of *A. boucheti*, the clustering of individuals according to axis 2 (sex) suggested that the group of unsexed individuals was composed by 11 females and 8 males (overall: 23 females, 18 males, *r* = 0.44, *p* = 0.53).

Finally, there was no visible effect of sex (that is, considering males versus females combined with morphological hermaphrodites) in the PCA of *A. kojimai* genotypes (Fig. 3C). With this species, the only obvious structuring factor was the main West-East geographic separation of sampling locations (PCA axis 1, 1.2% of the total variance). Importantly, this pattern was left unchanged when considering three groups (males, females, and morphological hermaphrodites identified from histology) rather than two (Fig. S5). Only one morphological hermaphrodite was identified in the Western region, but this is due to the fact that histological analyses concentrated on Eastern sites.

Within the Eastern region, no genetic structure was apparent between morphological hermaphrodites, females, and males (Fig. S5).

To quantify the role of sex in the distribution of genetic variation in the three *Alviniconcha* species, we estimated population genetics parameters considering males and females (or females combined with morphological hermaphrodites in the case of *A. kojimai*) for each species (Table 1). The genetic differentiation between sexes was low but significantly different from zero in all three species. In agreement with results from PCA, it was strongest in *A. boucheti* (*F*_ST_ = 0.0105, *p*-value < 0.001), followed by *A. strummeri* (*F*_ST_ = 0.0024, *p*-value < 0.001) and *A. kojimai* (*F*_ST_ = 0.0002, *p*-value = 0.037), reflecting minute differences in gene diversity within sexes versus overall individuals (Table 1). In species *A. kojimai* this genetic differentiation between sexes no longer existed when calculated with a reduced dataset composed by individuals identified as males, females, and morphological hermaphrodites from histology exclusively (see supplementary material text and Table S2).

### Sex determination system

In *A. boucheti*, 573 SNPs on 242 RAD-loci were strictly homozygous in females (*n* = 46), which all shared the same genotype, and heterozygous in males (*n* = 51, Figure 4a, top-left corner), pointing towards a XY genetic system of sex determination where each of these SNPs has one allele fixed on the X chromosome and the other fixed on the Y. By contrast, no SNP was found to be indicative of a ZW system. Furthermore, a large number of additional SNPs showed a combination of strong *H*_O,m_ and low *H*_O,f_, in agreement with the hypothesis of a XY system where SNPs will be linked to the sex chromosome but located in a still recombining region (see next section below).

**Figure 4:**
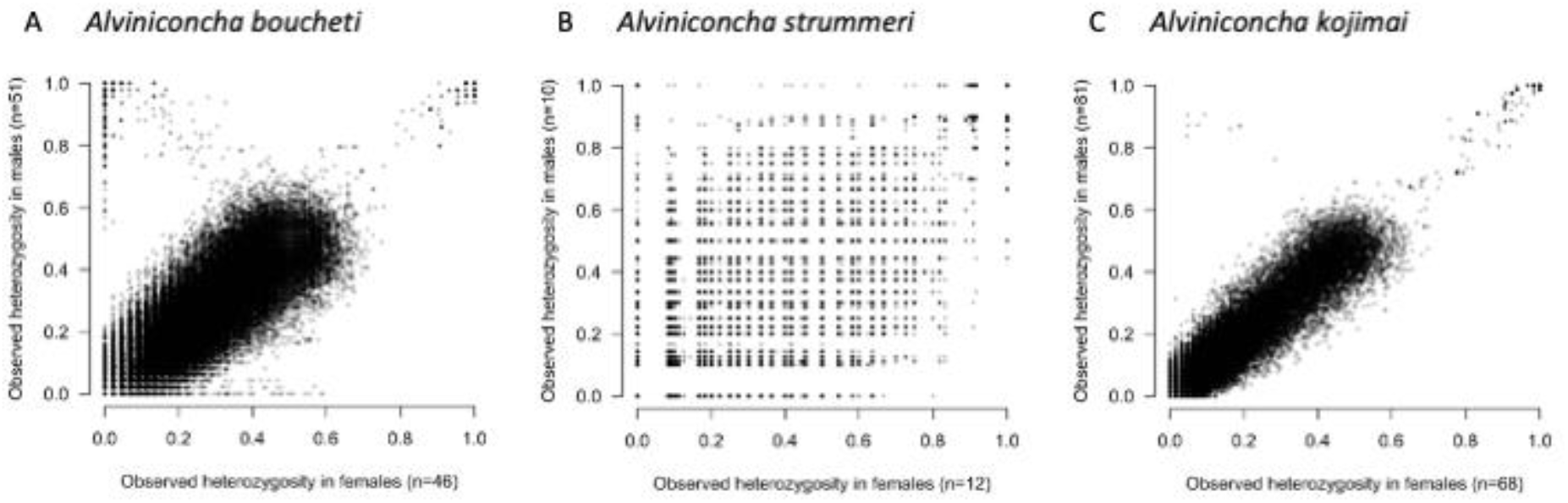
Locus by locus female and male-specific observed heterozygosity *H*_O,f_ and *H*_O,m_ in three *Alviniconcha* species. A) *A. boucheti*: 163 117 SNPs (46 females and 51 males). In the top-left corner, 573 SNPs (on 242 RAD) have *H*_O,f_ = 0 and *H*_O,m_ = 1, hence compatible with position of these SNPs on XY sex chromosomes with distinct alleles fixed on X and Y; B) *A. strummeri*: 36 568 SNPs (12 females and 10 males). In the top-left corner, 92 SNPs (on 55 RAD) have *H*_O,f_ = 0 and *H*_O,m_ = 1. Note that sample size was low in this species (22 sexed individuals), resulting in noisy estimates of *H*_O,f_ and *H*_O,m_; C) *A. kojimai*: 70 122 SNPs (68 hermaphrodites/females and 81 males). In the top-left corner, nine SNPs (on 8 RAD) showed strong heterozygosity in males but low heterozygosity in hermaphrodites/females.

For *A. strummeri*, although we had much fewer sexed individuals (12 females and 10 males) and thus a higher error rate associated with *H*_O,m_ and *H*_O,f_ at each locus, we found that 92 SNPs (on 55 RAD-loci) were strictly homozygous in females and heterozygous in males (Figure 4b, top-left corner), again indicating an XY genetic system of sex determination.

Surprisingly, in the species *A. kojimai*, where at least a third of the individuals have a mixture of male and female gonadal tissues, we still found 9 SNPs (on 8 RAD-loci) with strong *H*_O_ in males but low in hermaphrodites/females (top-left corner of Figure 4c). This pattern, which remained consistent when separating males, females, and morphological hermaphrodites into three groups (Fig. S7), again pointed towards a XY-like system where males are XY and morphological hermaphrodites / females are XX (see Discussion). However, this sex determination was supported by a very small number of SNPs (9 out of 70 122, about 0.13 %) and contrary to the two other species, none of these SNPs were strictly homozygous in females and heterozygous in males.

In all three species, a number of SNPs had simultaneously high *H*_O,m_ and *H*_O,f_ values (top-right corner of Figs. 4a-c) possibly indicative of paralogous loci that were wrongly assembled together, or, perhaps less likely, sites under strong balancing selection.

### Sex-linked loci

In *A. boucheti*, 1 011 SNPs (on 373 RAD-loci) had a combination of low *F*_IS,m_ and strong *F*_ST_ compatible with XY features that were never obtained by simulation of autosomal segregation (Figs. 5A-B). This represented ca. 0.7% of the set of SNPs for which *F*_IS,m_ and *F*_ST_ could be calculated (blue dots in Fig. 5B, *n* = 142 230).

**Figure 5:**
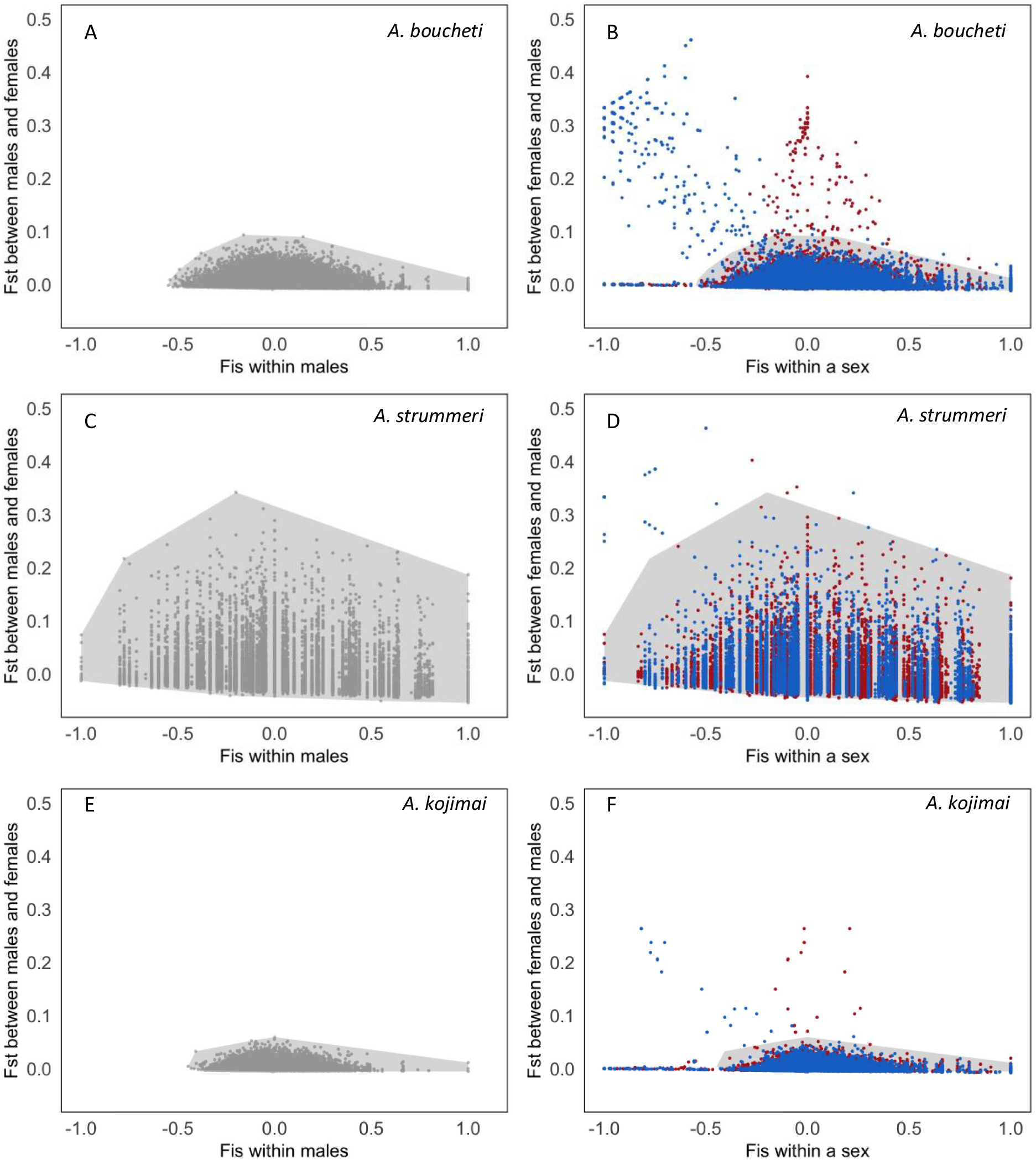
Simulated and observed *F*_ST_ values between males and females and sex-specific *F*_IS_ for three *Alviniconcha* species (top row: *A. boucheti*, middle: *A. strummeri*, bottom: *A. kojimai*). Panels A, C, and E show simulated values obtained for autosomal SNPs with a range of expected allelic frequencies (uniform between 0 and 1) and using real male and female sample sizes from each species (table 1). The grey polygon is a convex envelope that covers all possible values obtained by simulation for an autosomal locus. Panels B and D show observed values for male *F*_IS_ (in blue) or female *F*_IS_ (in red) and male/female *F*_ST_ (i.e., each SNP is represented once in blue and once in red according to each gender).

Similarly, in *A. strummeri*, 111 SNPs (on 64 RAD-loci) out of 32 397 SNPs had *F*_IS,m_ and *F*_ST_ that were never obtained by simulation of autosomal SNPs (Figs. 5C-D). These SNPs represented 0.34% of the markers analysed here (blue dots in Fig. 5F, *n* = 32 397), but the power for detecting sex-linked SNPs was very reduced for *A. strummeri* because only 22 individuals were sexed, and this number is often even slightly lower for each specific locus (due to locus-specific missing data).

Panel F is similar but, in this case, red symbols refer to morphological hermaphrodites and females, and *F*_ST_ was calculated between males and morphological hermaphrodites/females. Any SNP outside of the grey polygon has properties that were never obtained by neutral autosomal simulations.

Finally, in *A. kojimai*, 16 SNPs (on 14 RAD-loci) had *F*_IS,m_ and *F*_ST_ values that were never obtained by simulation of autosomal SNPs (Figs. 5E-F). This represented 0.025% of the set of SNPs for which *F*_IS,m_ and *F*_ST_ could be calculated (*n* = 64 310).

In line with the expectations for XY sex chromosomes for the three species, all SNPs showing strong differentiation between sexes (*F*_ST_) tended to have negative *F*_IS_ values in males (as stated above) and null *F*_IS_ in females (shown in red in Fig. 5).

### Loci present only on the Y chromosome

In species *A. boucheti*, we found 89 SNPs (on 40 RAD loci) that were absent from all females while genotyped in at least 10 males. These RAD loci may be located in genomic regions that are specific to the Y chromosome, or they may be homologous but too divergent between X and Y to be recognized as such by our analyses. In contrast, no such loci were found in *A. strummeri* and *A. kojimai*.

### Synteny of sex-linked loci in the three Alviniconcha species

Setting aside the Y-specific *A. boucheti* loci, a reciprocal BLASTn of sex-linked loci between pairs of species revealed that 28 of them were common to at least two species (including 3 RAD loci common to the three species, Fig. 6A). In particular, 10 of the 14 RAD loci identified as sex-linked in *A. kojimai* were also found to be sex-linked in *A. boucheti* (*n* = 4), *A. strummeri* (*n* = 3), or both (*n* = 3). By contrast, a large proportion of loci were found to be sex-linked in *A. boucheti* only (n = 348), or *A. strummeri* only (n= 40), which may be due to differences in sample sizes and statistical power, or the length of sex chromosomes (see discussion).

**Figure 6:**
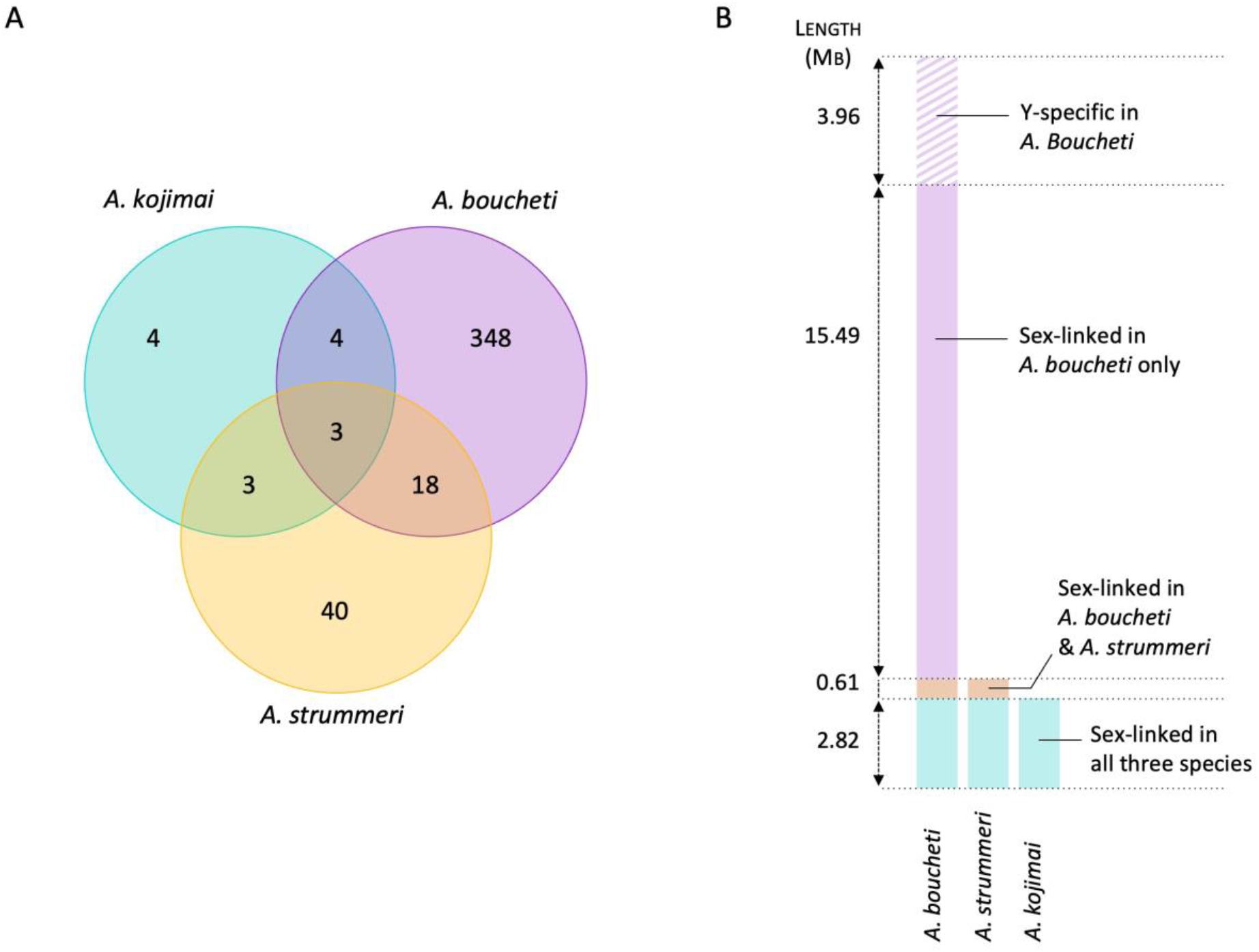
A) Distribution of sex-linked RAD-loci shared by *A. boucheti* (97 individuals, 373 sex-linked RAD loci), *A. strummeri* (22 ind., 64 loci) and *A. kojimai* (149 individuals, 14 loci). B): total length and interspecies distribution of genomic scaffolds found to be sex-linked in each species, using *A. marisindica* reference genome (Yang *et al*., 2022). These sizes are indicative and should not be taken as face values since they depend on the contiguity of the genome assembly used as a reference (especially in *A. boucheti*) and the unknown frequency of repeated elements on sex chromosomes.

Eleven of the 14 sex-linked RAD-loci identified in *A. kojimai* were retrieved by blastn on 5 scaffolds of the *A. marisindica* genome assembly (the remaining three loci also mapped on these scaffolds, but with a size coverage of less than 95%). The total length of these scaffolds amounted to 2.8 Mb (Fig. 6B). Most importantly, these 5 scaffolds were also found to include sex-linked RAD-loci for the two other species: *A. boucheti* (67 RAD-loci out of 373) and *A. strummeri* (15 RAD-loci out of 64).

Out of the 64 sex-linked loci identified in *A. strummeri*, 20 returned a significant hit on the *A. marisindica* genome. They were distributed on 8 scaffolds (total length 3.4 Mb) that included the 5 scaffolds identified for *kojimai* loci. Each of these eight scaffolds also included 1 to 34 RAD loci found to be sex-linked in *A. boucheti*.

Finally, 259 of the 373 sex-linked RAD loci identified in *A. boucheti* segregated on 41 scaffolds, with a total length of 21.7 Mb. In addition, 21 Y-specific RAD loci were found on 18 scaffolds, seven of which seemed to bear only Y-specific loci (concatenated length 3.96 Mb).

### Phylogeny of sex-linked RAD haplotypes

The three RAD loci that were sequenced and identified as being sex-linked in all three species (Fig. 6A) were located on two of the five sex-linked scaffolds shared by the three species (Fig. 6B). The phylogeny of each of these loci shows that haplotypes cluster primarily by species, and then by gametologs within each species (Fig. 7). These genealogies reflect the fact that the haplotypes observed on a given sex chromosome (e.g. X) were always more similar to the haplotypes observed on the opposite sex chromosome (Y) within the same species rather than to the haplotypes observed on the same gametolog (X) in other species. Nevertheless, in *A. boucheti* and *A. strummeri*, all copies of the Y chromosome shared a single haplotype (per species) that was different from all haplotypes observed on the X chromosomes (with a single exception on locus 3 in *A. boucheti*, Fig. 7). In contrast, in *A. kojimai*, the X and Y chromosomes shared one to three haplotypes per locus.

**Figure 7:**
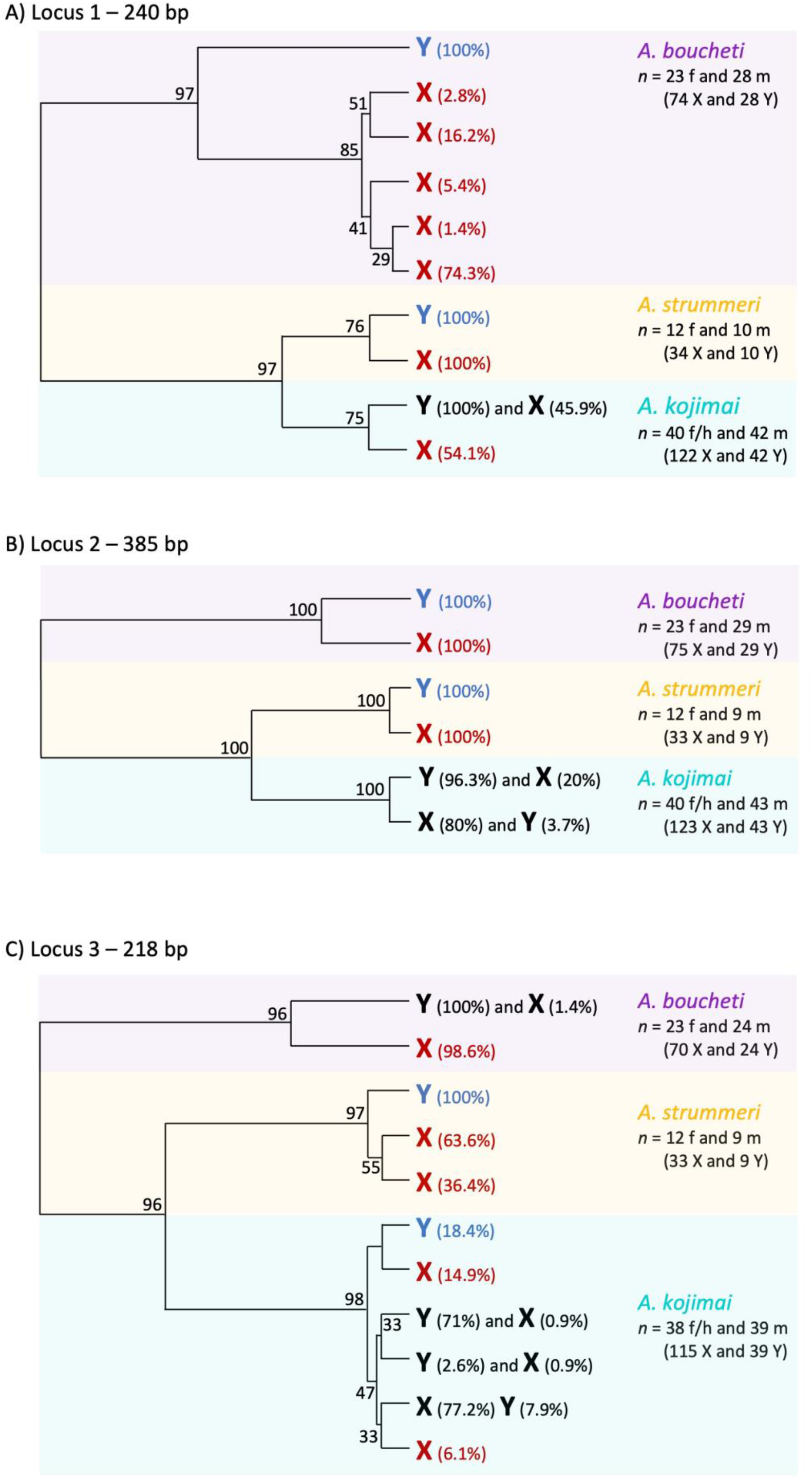
Phylogeny of three RAD-seq loci located on the sex chromosomes of *Alviniconcha boucheti*, *A. strummeri*, and *A. kojimai*. Each tree was constructed using 145 to 155 individuals with phased haploid sequences (i.e. 290 to 310 X or Y haploid sequences per locus). The red symbol “X” and the blue “Y” indicate that the haplotype was found only on X or Y chromosomes, respectively. The percentages indicate what proportion of sequences corresponded to a particular haplotype (e.g. the first X in the top panel corresponds to a haplotype that was observed twice, only in *A. boucheti*, and each time on an X chromosome, which means that it represented 2/74 = 2.8% of all X sequences in the species *A. boucheti*. The same logic applies to the blue “Y” symbols. The black color is used for haplotypes that were found on both sex chromosomes. For example, in panel A, all Y chromosomes of *A. kojimai* carried the same haplotype, and this haplotype was also found on almost half of the X chromosomes.

## Discussion

Anatomical and histological observations of gonads, combined with analyses of sex-linked genetic variation, informed us about the sexual and sex determination systems of poorly known deep-sea invertebrates. In particular, the observation of hermaphrodites or intersex individuals in at least one species of *Alviniconcha* gastropods suggests that this genus provides an opportunity to study sexual systems transition and the role of the landscape of recombination on transitions.

### Sexual and sex determination systems in Alviniconcha gastropods

We found that two species (*A. boucheti* and *A. strummeri*) are gonochoric with an XY genetic system of sex determination, and balanced sex-ratio. This conclusion is based on unequivocal observations of gonad cross sections (showing male-or female gonadic tissue), and sex-specific distribution of the genetic variation (1011 and 111 sex-linked SNPs in *A. boucheti* and *A. strummeri*, respectively, including in particular 573 and 92 SNPs with alleles differentially fixed on X and Y chromosomes).

The third species, *A. kojimai*, presented a strikingly different pattern. Close examination of histological cross-sections revealed that approximately 75% of the individuals that had been classified on board the ship as female based on external observations of the gonad actually had a mixture of female and male gonadal tissue (Fig. 2). Interestingly, however, while no SNP was found to be strictly heterozygous in *A. kojimai* males, nine markers located on seven RAD loci showed observed heterozygosity values (high in males, low in females) that were clearly outside the pattern defined by the other 70 113 SNPs (Fig. 4C), again suggesting an XY system. Importantly, the analyses comparing males vs. females + morphological hermaphrodites were confirmed by the analyses restricted to those individuals for which sex was determined through histological observations: sex-linked outlier loci were detected when comparing males *vs* females or males vs morphological hermaphrodites, but not when comparing females vs hermaphrodites (Figs. S7 and S8). Females and hermaphrodites therefore form a homogeneous sex group. Taken together, these results indicate that sex in *A. kojimai* is at least partially determined genetically, with XY individuals developing into males and XX individuals developing into females or intersex individuals with varying amounts of male gonadal tissue.

### A single pair of XY chromosomes across the three species, but different recombinational landscapes

Reciprocal blastn of sex-linked loci for each pair of species revealed that three RAD loci were found to be sex-linked in the three species, and 25 more were identified as sex-linked in two species (Fig. 6A). In addition, the five scaffolds of the *A. marisindica* reference genome containing all 14 *A. kojimai* sex-linked RAD did also include sex-linked loci from the two other species (Fig. 6B). These results demonstrate that the three species share this sex-linked genomic region, which contains or is close to the region that triggers sex determination in the three species, on the same pair of XY chromosomes. This pair of sex chromosomes is presumably ancestral to all species, but we cannot rule out the possibility that it became sex-linked independently in each species.

It is difficult to estimate the size of the sex-linked region in each species because the number of sex-linked loci identified depends on sample size (number of sexed individuals, which is an issue for *A. strummeri* in particular), methodology (conservative threshold set by our simulations to decide whether a locus is sex-linked), XY polymorphism (itself depending e.g. on the species effective population size and structure), and imprecisions introduced by repeated elements.

Our reasoning in analyzing sex-linked genetic variation is based on the idea that the increase in genetic differentiation between homologous sex chromosomes is due to a reduction in recombination, or even its complete cessation in the case of loci that appear to be differentially fixed on the X and Y chromosomes (e.g. Berset-Brändli *et al*., 2006; Brelsford *et al*., 2017). Yet one must keep in mind that sex chromosomes may contain repeated elements, which could create spurious RAD-loci identified as homologous sequences. Such repeated elements can accumulate mutations that would be wrongly identified as heterozygous SNPs. Tandem replications of non-coding DNA, and in particular transposable elements can accumulate as a direct consequence of recombination arrest and may end up representing a large fraction of Y chromosomes (Beukeboom & Perrin, 2014; Charlesworth, 2021). For instance, some of the SNPs that we identified in *A. boucheti* and *A. strummeri* as having one allele fixed on the X and another one fixed on the Y could in fact have the two alleles on distinct repeated sequences on the Y. Nonetheless, the conclusion remains the same: these loci are a good indicator of XY genetic sex determination, but they should not be automatically seen as simple homologous markers present in a single copy (the results shown in figure 4 do in fact suggest that there are a number of autosomal paralogs, identified as loci with simultaneously high male and female heterozygosities).

A first approach to the size of sex-linked regions is simply to relate the proportion of sex-linked loci to the total size of the genome in the two species where sample size provided enough power. In *A. boucheti*, 0.7% of the SNPs were identified as sex-linked, while this figure dropped to 0.025% in *A. kojimai*. The total genome size was previously estimated for two *Alviniconcha* species: 0.92 Gb for an unidentified specimen from one of our species (formerly called *A. hessleri* before taxonomic revision) estimated by flow cytometry (Bonnivard *et al*., 2009), and 0.83 Gb for the genome assembly ASM1885773v1 of *A. marisindica* (Yang *et al*., 2022). If *A. boucheti* and *A. kojimai* have a genome of about 0.9 Gb then 0.7% and 0.025% of this genome represent sex-linked regions of 6.3 Mb for *A. boucheti* and 2.25 Mb for *A. kojimai*, respectively.

In a second approach, we summed the lengths of the *A. marisindica* genome scaffolds found to harbor sex-linked loci. The five such scaffolds identified for *A. kojimai* (and that also host a large number of sex-linked loci in the other two species) have a total length of 2.82 Mb (Fig. 6B). This result depends of course on the continuity of the reference genome used, but it is quite close to the estimate reported above. By contrast, in *A. boucheti* we found many more scaffolds with sex-linked loci (total size 21.7 Mb), and we also found an additional genomic region that seems to be specific to the Y chromosome and absent from the X (total size 3.96 Mb). These values for *A. boucheti* should be interpreted as rough, minimal estimates because we used rather conservative thresholds for mapping loci on the reference genome, and most importantly we kept only the best hit, ignoring multiple mappings potentially due to repeated elements. The true size of the sex-linked region in *A. boucheti* might be larger than reported here. And the same caveat applies to *A. strummeri*, where the power to identify sex-linkage might have been impeded by sample size.

A precise estimation of the size and synteny of sex-linked genomic regions in the different *Alviniconcha* species will require full genome sequences. Yet the conclusion that the sex-linked region is much larger in *A. boucheti* than *A. kojimai* is a robust one. One can hypothesize that sex-chromosomes are smaller in *A. kojimai*, and/or that in this species, recombination occurs on a much larger portion of the X and Y chromosomes (a hypothesis that would fit with the fact that we did not find any fully heterozygous SNP in males in this species). It is also possible that there was a recent breakdown of the non-recombining region in *A. kojimai*.

The role of recombination in the evolution of sex chromosomes is supported by the phylogeny constructed for three sex-linked RAD-loci (Fig. 7): although the three *Alviniconcha* species share the same pair of XY chromosomes, X haplotypes were always more similar to Y haplotypes within a given species than they were to allospecific X haplotypes. It is possible that the same pair of autosomes became sex-linked independently within each species at different points in time, but a more likely hypothesis is ancestral X and Y chromosomes have experienced recombination in the genomic region harboring our three RAD loci within each species since speciation events (estimated by Breusing *et al*., 2020 to date 48 Mya for the split leading to *A. boucheti*, and 25 Mya for the split leading to *A. strummeri* and *A. kojimai*).

On the other hand, recombination must have been rare enough to allow X and Y haplotypes to diverge within each species to the extent that they became differentially fixed (or nearly so) in *A. boucheti* and *A. strummeri* (and branch lengths in figure 7 show that the genetic distance between X and Y was repeatedly much higher in *A. boucheti* at all three loci). In contrast, in *A. kojimai*, most haplotypes are shared between X and Y chromosomes, showing that our three loci are located in a genomic region where recombination may have limited X-Y divergence and rejuvenated the Y more than in *A. boucheti*. Along the same line, branch lengths for locus 1 show that X-Y divergence in *A. kojimai* (and even *A. strummeri*) is comparable to that between *A. boucheti* X haplotypes. This means that, at this particular locus, X-Y divergence did not build up in *A. kojimai* and *A. strummeri* as it did in *A. boucheti*. Given the surprising sexual system of *A. kojimai*, it will be interesting to investigate further the relationship between XY structure, recombination, and sex determination in this species, in relation to all other species of the genus.

### What is the sexual system of Alviniconcha kojimai, and how is sex determined in this species?

As for the two other Alviniconcha species, all our observations point towards a XY system of sex determination in *A. kojimai*, with XX individuals developing a mixture of male and female tissues in variable proportion. We have yet to determine if the mixed gonadal tissues allow these individuals to reproduce using both sex functions, and, if so, simultaneously or not, and whether selfing is possible. Here we analysed a single gonad cross section per individual, and further investigations are therefore needed to quantify these proportions precisely, and in particular, determine if some females that do not show any male tissue are indeed devoid of male tissue (11 such individuals identified here using a single gonad section, i.e. 12% of all individuals used in histological analysis). All this information is necessary to understand the current sexual system of *A. kojimai*, and to understand whether it may be stable or represent a transitional stage towards gonochorism or hermaphroditism.

If the male tissue observed in the female gonads is non-functional, then *A. kojimai* is effectively a gonochoric species. The balanced sex-ratio observed from histological analyses (48 males, 42 females or hermaphrodites, *r*=0.53, binomial test *p*=0.6) is compatible with this hypothesis. However, the fitness costs of non-functional male tissue in female gonads should generate a strong opportunity for selection against alleles conducive to intersex phenotypes. Surprisingly, it seems that many gonochoristic species appear to contain individuals that have both male and female reproductive traits, but with only one type being functional (i.e. a single type of functional gamete produced and used for reproduction). In the perspective of the evolution of sexual systems, this condition has been discussed mostly in the botanical literature (where it is called cryptic dioecy, e.g. Charlesworth, 1984; Mayer & Charlesworth, 1991). In the animal biology literature, this topic has been more intensively examined in the perspective of the anthropogenic stressors possibly causing the development of reproductive tissue typically seen as abnormal (reviewed in Bahamonde *et al*., 2013; Grilo & Rosa, 2017). Some gonochoristic animal species appear to contain individuals called intersex that present both male and female traits. This is particularly frequent in fish and molluscs, where intersex refers more specifically to “individuals possessing oocytes or distinct stages of spermatogonia, at varying degrees of development, within the normal gonad of the opposite gender (i.e. spermatocytes in the ovary or oocytes in the testis)” (Grilo & Rosa, 2017). Intersex was shown to be exacerbated by environmental contaminants that can disrupt sex differentiation, and thus should be detrimental for the species. However, exogenous factors may not be necessarily required, and the frequency of “naturally” occurring intersex individuals remains an open question (at the exception perhaps of the case of imposex, a strong masculinization of marine gastropod females caused by endocrine disrupting chemicals, Gibbs & Bryan, 1996; De Wolf *et al*., 2001). In all cases except imposex, these species are in fact functionally gonochoristic, but they are interesting as they may nonetheless represent intermediate steps in transitions between fully functional sexual systems (e.g. in angiosperms, Mayer & Charlesworth, 1991). Here however, the evolutionary success of *A. kojimai* in colonizing nearly all hydrothermal vents of four back-arc basins and the volcanic arc of Wallis & Futuna despite a high ratio of morphologically hermaphroditic individuals found in every population should indicate that such a “transitional” stage has been favoured for quite a long time (at least a few thousands generations, Castel *et al*., 2022).

Considering the anatomical observations only, another possibility could have been that the intersex individuals were undergoing sex change (sequential hermaphroditism). For instance, in a recent study of *Gigantidas haimaensis* mussels from cold seep ecosystems in the South China Sea, Shi *et al*. (2022) found that male gonads contained oocytes (while female gonads did not contain spermatocytes). Combining proteomic and transcriptomic analyses with their histological observations, the authors hypothesized that these males were intersex individuals undergoing sex change, a strategy observed in other Bathymodiolinae (Laming *et al*., 2018) possibly favoured by the ephemeral and patchy distribution of deep sea vent and cold seep habitats. But in the case of *A. kojimai* this hypothesis is refuted by our inference of XY genetic sex determination.

Finally, the male tissue observed within female gonads may be functional. That would make *A. kojimai* an androdioecious species, where males coexist with functional hermaphrodites that effectively gain reproductive success through both their male and female functions (Charlesworth, 1984; Pannell, 2002). One may even consider trioecy if some *A. kojimai* females really do not contain any male tissue at all within their gonads (a possibility that remains to be investigated). However, there are in fact very few species with such mixed systems: Weeks (2012) listed only 115 androdioecious animals (none reported in molluscs). The presence of male tissue within hermaphrodites would be only beneficial if the sperm is used for selfing or cross-interbreeding via sperm storage when the number of individuals is restricted to found a new population: a situation possibly met when geo-tectonic rearrangements or eruptive events destroy most of the vent communities in a given region (see the recent eruption of the submarine volcano Hunga Tonga – Hunga Ha’apai in December 2021, Gupta *et al*., 2022). Androdioecy could explain the success of *A. kojimai*, which we found to have a larger range and a higher abundance than its syntopic congener *A. strummeri* (Castel *et al*., 2022).

Mixed sexual systems are particularly relevant to understand the evolution of sexual systems, and may in some cases represent intermediate stages between hermaphroditism and gonochorism (Charlesworth, 2006; Pannell & Jordan, 2022). Although the ultimate forces that drive transitions between gonochorism and hermaphroditism in animals are far from fully resolved, selection may arise from the costs and benefits of factors such as reproductive assurance, inbreeding, sexual specialization, and plasticity in sex allocation (Weeks, 2012; Leonard, 2018; Pannell & Jordan, 2022). Gonochorism is thought to be the ancestral sexual system of gastropods (Collin, 2013) but molluscs have very diversified sexual systems relatively to other major animal clades, with much flexibility in sex determination and sex differentiation (Collin, 2013; Leonard, 2018). Collin (2013) mentions in particular that intersex individuals were observed in many molluscan taxa, sometimes in non-negligible numbers, and that these species deserve further investigations to ascertain their sexual system. Extending the analyses presented here to sexual anatomy and sex-linked genomic patterns of the six species of *Alviniconcha* gastropods (Johnson *et al*., 2015) should prove interesting with respect to the evolution of sexual systems and sex determination.

## Acknowledgements

We thank the crew of the RV L’Atalante and the team of the ROV Victor6000 who participated in the Chubacarc 2019 expedition. We thank P-A. Gagnaire, who suggested that we look at the sex of individuals to understand the observed genetic structure of *A. boucheti*, and we are grateful to Mathieu Gautier for insightful discussions. Alan Le Moan provided key advices with genetic principal component analyses. This work benefited from access to the Genomer molecular biology platform, member of BioGenOuest and EMBRC-France, and the bioinformatics platform ABiMS of the Station Biologique de Roscoff.

## Funding

This study was supported by the French Oceanographic Fleet program and the ANR CERBERUS (ANR-17-CE02-0003 to S. H.), and the ARED program of the Region Bretagne.

## Supplementary information

Supplementary material is available on biorxiv together with the article (doi: 10.1101/2023.04.11.536409).

## Data, script, and code availability

All ddRAD-sequences and associated metadata are available on NCBI within the BioProject PRJNA768636 (accessible at https://www.ncbi.nlm.nih.gov/bioproject/PRJNA768636). R-scripts and additional data are deposited on zenodo.org (DOI 10.5281/zenodo.10078642), accessible at https://doi.org/10.5281/zenodo.10078642.

## Conflict of interest disclosure

The authors declare they have no conflict of interest relating to the content of this article. T. Broquet is a recommender for PCI Evolutionary Biology.

## Notes

https://doi.org/10.5281/zenodo.10078642

### Competing Interest Statement

The authors have declared no competing interest.

### Summary of Updates

Following recommendation of this preprint by Peer Community in Evolutionary Biology, a PCI badge with a link to the recommendation was added on the front page.

