## Supplementary material for "Genetic sex determination in three closely related hydrothermal vent gastropods, including one species with intersex individuals": Suplementary material

### Sampling

Table S1 - Number of females, males, and unsexed individuals per sampling location. In *Alviniconcha kojimai*, column "F or H" indicates the number of females (no male tissue visible in the histological section) and morphological hermaphrodites with varying proportions of male tissue. The sex indicated in this table is the sex observed from histological analyses (most reliable method) for 166 individuals, and the sex observed from the external colour of the gonad for the remaining 105 individuals. All numbers refer to the samples that passed all filters.

| Basin or volcanic arc | Vent field | *A. boucheti* | | | *A. strummeri* | | | *A. kojimai* | | |
|---|---|---|---|---|---|---|---|---|---|---|
| | | F | M | U | F | M | U | F or H | M | U |
| Manus | Pacmanus | 18 | 12 | 43 | - | - | - | - | - | - |
| | Susu | 1 | 2 | 45 | - | - | - | 6 | 9 | 31 |
| Woodlark | La Scala | 9 | 10 | 1 | - | - | - | 2 | 3 | 18 |
| North Fidji | Phoenix | - | - | - | 3 | 3 | - | 11 | 14 | 6 |
| Futuna | Fati-Ufu | 2 | 2 | - | 1 | 2 | 1 | 16 | 13 | 6 |
| | Fatu-Kapa | - | - | - | - | - | - | 6 | 11 | - |
| Lau | Mangatolo | 8 | 9 | - | - | - | - | 9 | 9 | - |
| | Tow Cam | 3 | 6 | 9 | - | - | - | 3 | 7 | 23 |
| | ABE | 6 | 12 | 1 | - | 1 | - | 1 | 1 | - |
| | Tu'i Malila | - | - | - | 8 | 4 | 18 | 14 | 14 | 13 |
| *Total* | | *47* | *53* | *99* | *12* | *10* | *19* | *68* | *81* | *97* |

### Selection of parameters for ddRAD-seq data analysis

Within each species, assembly parameters were chosen based on a pilot study of a subset of samples (*A. boucheti*: 28 samples, *A. kojimai*: 30 samples, *A. strummeri*: 46 samples). These sample subsets included triplicates of individuals for two species: six *A. boucheti* and six *A. kojimai* were included three times independently in the ddRAD-seq library preparation, starting from a unique DNA extract, therefore allowing us to estimate genotyping errors for these two species. For each pilot study, we varied one Stacks core parameter at a time (by steps of one unit: ustack *m* [3–7], *M* [2–8], and cstack *n* [2–12], while the others remained at their default values, as suggested by Mastretta-Yanes et al. (2015). All analyses of these pilot studies used the following set of constant conditions: max_locus_stacks was set to 3, the minimum proportion of individuals sharing a locus was set to *r* = 0.8, and the minor allele count set to MAC = 4. The SNPs identified by Stacks were then further filtered in R version 4.1.1 (R Core Team, 2021) in order to keep the SNPs present in at least 90% of the individuals and keep only the individuals which were genotyped at least at 80% of the total number of SNPs, except in *A. strummeri*, for which we had less individuals and thus allowed a little more missing data per individual (25%).

The final set of parameters was chosen such as to maximise the number of single nucleotide polymorphism sites (SNPs) retained after filtering while maintaining a minimum genotyping error rate (measured as the frequency of differences in the three genotypes observed at each SNP and each replicate). As *A. strummeri* had no replicates in our dataset, for this species the choice of parameters was based only on the number of SNPs retained. The results of these pilot studies are presented in figures S1-3. The optimal values that minimize the genotype differences between replicates and maximize the amount of data recovered were identified as $m$=6 (all species), and $n$=$M$ set to 4, 8, and 10 for *A. boucheti*, *A. strummeri*, and *A. kojimai*, respectively. We allowed high values for $M$ and $n$ parameters to account for the potential occurrence of divergent haplotypes within some individuals due to introgression between divergent species, particularly in *A. strummeri* (Castel *et al.*, 2022).

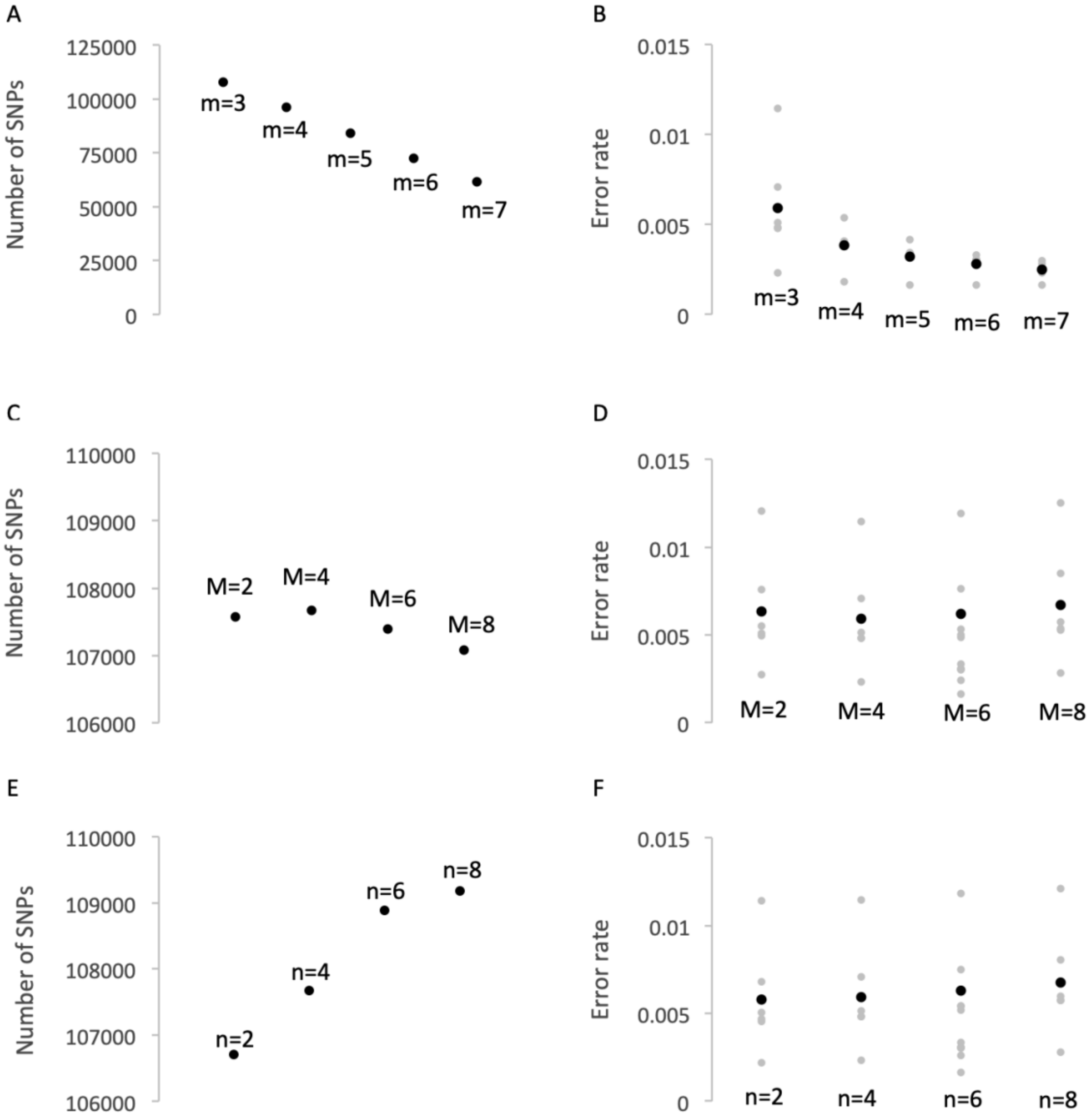


Fig. S1 - Selection of *Stacks* parameters for *Alviniconcha boucheti*. Number of single nucleotide polymorphisms (SNPs, left panels) and genotyping error rate (right panels) after data filtering for varying values of *Stacks* parameters *m*, *M*, and *n* using six *A. boucheti* individuals (3 replicates per individual). Note the limited range of the Y-axis in panels C and E.

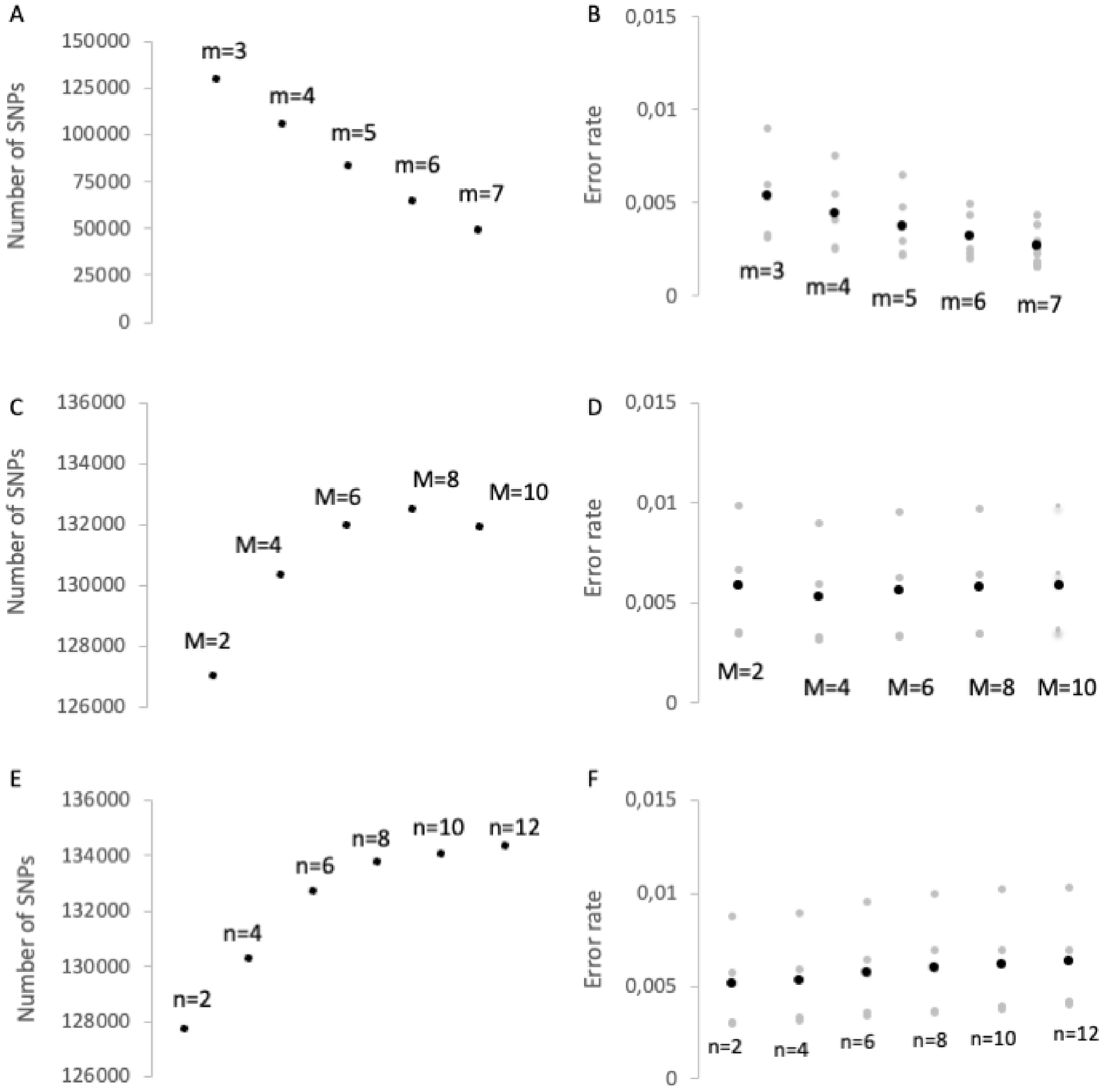


Fig. S2 - Selection of *Stacks* parameters for *Alviniconcha kojimai*. Number of single nucleotide polymorphisms (SNPs, left panels) and genotyping error rate (right panels) after data filtering for varying values of *Stacks* parameters *m*, *M*, and *n* using five *A. kojimai* individuals (3 replicates per individual). Note the limited range of the Y-axis in panels C and E.

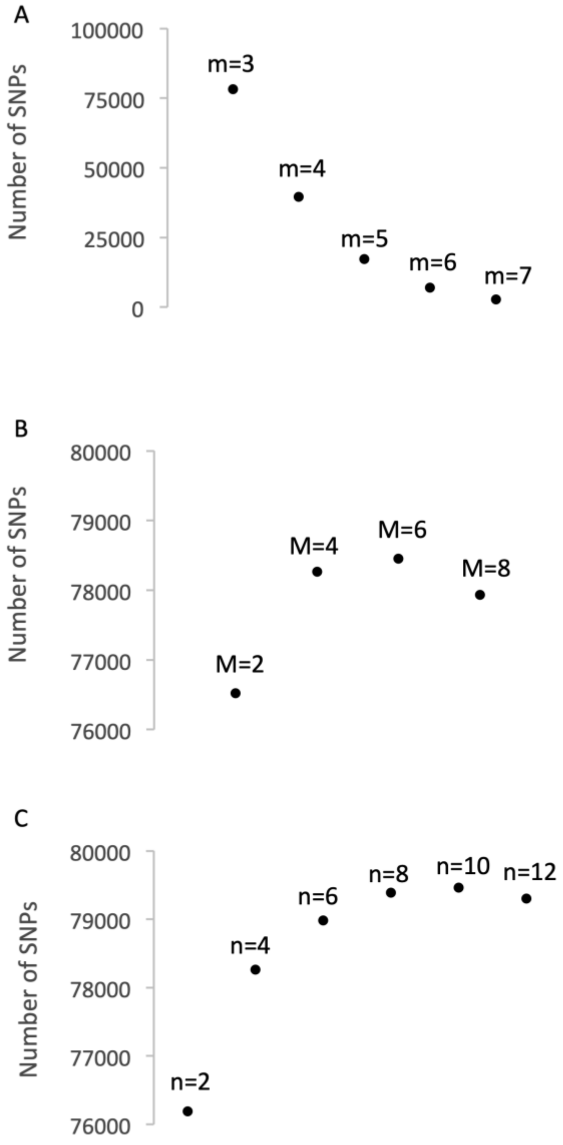


Fig. S3 - Selection of *Stacks* parameters for *Alviniconcha strummeri*. Number of single nucleotide polymorphisms (SNPs) after data filtering for varying values of *Stacks* parameters *m*, *M*, and *n*. Note the limited range of the Y-axis in panels B and C.

Table S2: Mean depth of coverage per SNP in the final dataset

| Species | mean per species | 2.5% quantile | 97.5% quantile |
|---|---|---|---|
| *A. boucheti* | 27.5 | 18.3 | 40.6 |
| *A. kojimai* | 25.3 | 19.1 | 35.0 |

| | | | |
|---|---|---|---|
| *A. strummeri* | 24.8 | 17.2 | 36.8 |

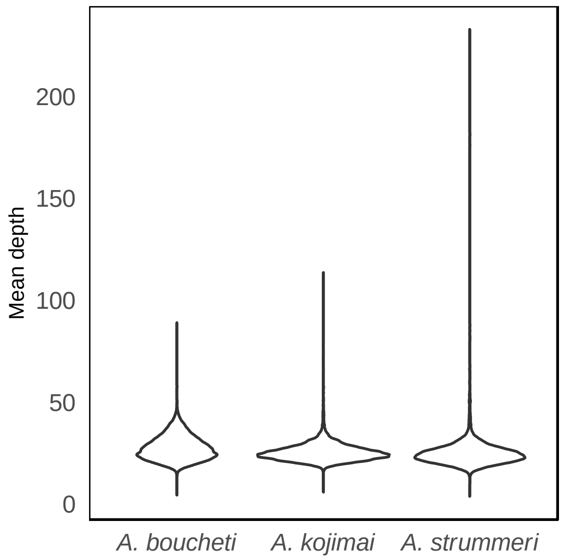


Figure S4: Distribution of the mean depth of coverage per SNP for each species.

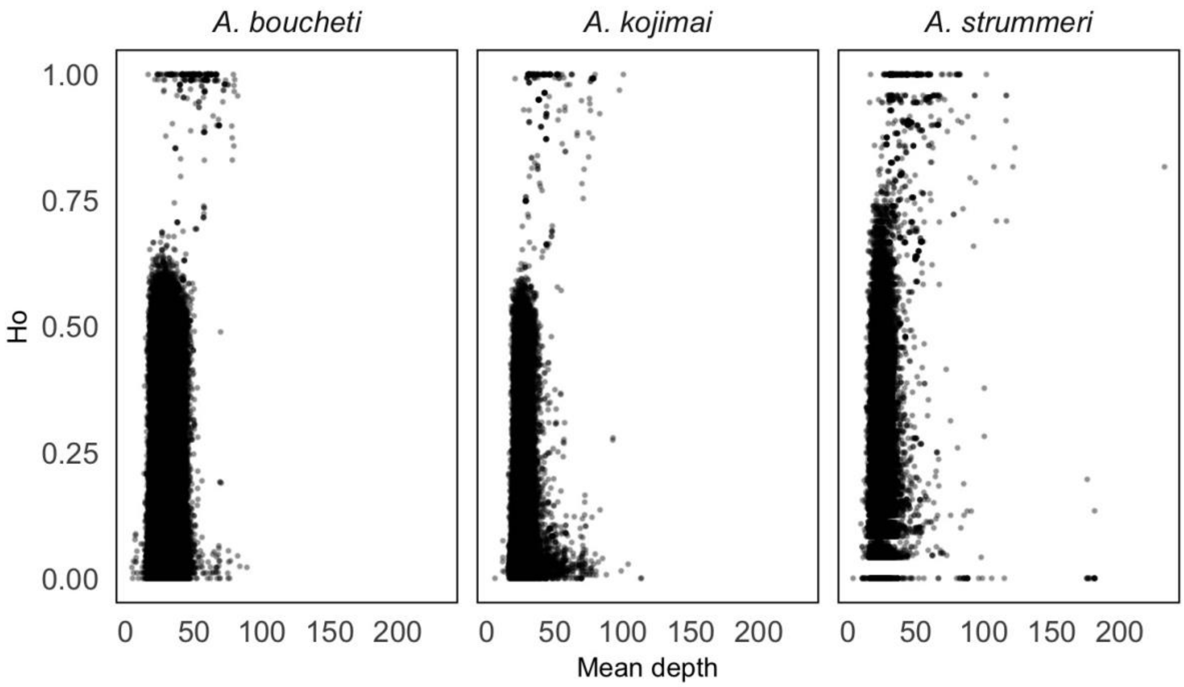


Figure S5: Observed heterozygosity as a function of mean depth of coverage per SNP.

**Genetic analyses performed separately for males, females, and morphological hermaphrodites in *A. kojimai*.**

The analysis of genetic diversity for *A. Kojimai* (presented in main text Table 1) was repeated using three sex classes: males, females, and hermaphrodites, identified using data from the histological analyses only. This represents 91 individuals, genotyped using 70122 SNPs. The global gene diversity $H_T$ was 0.1115 and $F_{ST}$ across sexes were as follows: $F_{ST,\ male\text{-}female}$ = -0.00034 ($p$-value = 0.9), $F_{ST,\ female\text{-}hermaphrodite}$ = -0.00016 ($p$-value = 0.7), $F_{ST,\ male\text{-}hermaphrodite}$ = 0.00014 ($p$-value = 0.127).

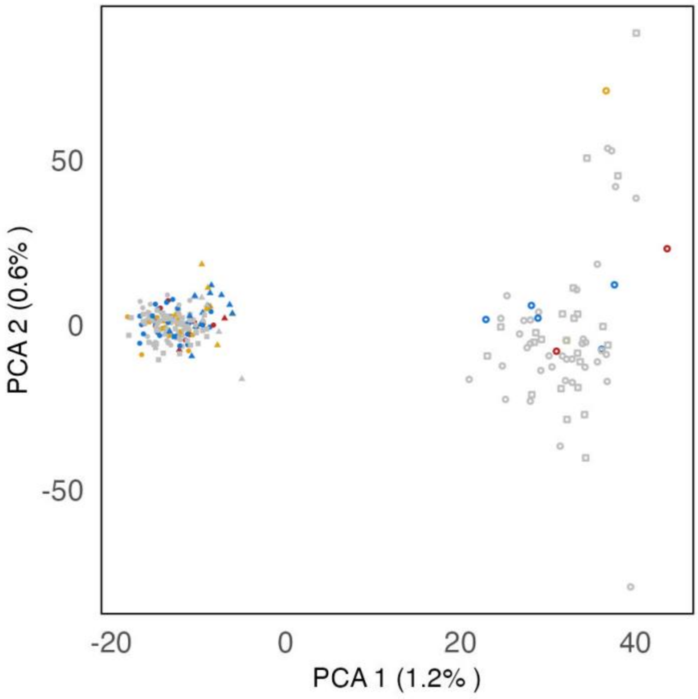


Figure S6: Principal components of SNP genotypes for *A. kojimai* (246 individuals, 70 122 snps). The first axis separates individuals according to their geographic origin: empty symbols correspond to samples from the Manus and Woodlark basins (north-west of the study area, see main text figure 1), solid symbols correspond to samples from the Futuna, North Fiji and Lau areas (South-East). There is no visible clustering by sex (orange symbols: morphologically androdioecious individuals, red symbols: individuals where only female gonadic tissue was observed, blue symbols: males).

Table S2 - Diversity indices for *A. kojimai* using histological analyses. $H_T$ : total gene diversity. $H_S$ : sex-specific gene diversity. $H_O$ : observed heterozygosity, and $F_{IS}$: heterozygosity deficiency.

| Sex | *n* | $H_s$ | $H_o$ | $F_{IS}$ |
|---|---|---|---|---|
| *Males* | 49 | 0.1119 | 0.1106 | 0.0114 |
| *Females* | 11 | 0.1118 | 0.1098 | 0.0177 |
| *Hermaphrodites* | 31 | 0.1108 | 0.1093 | 0.0136 |

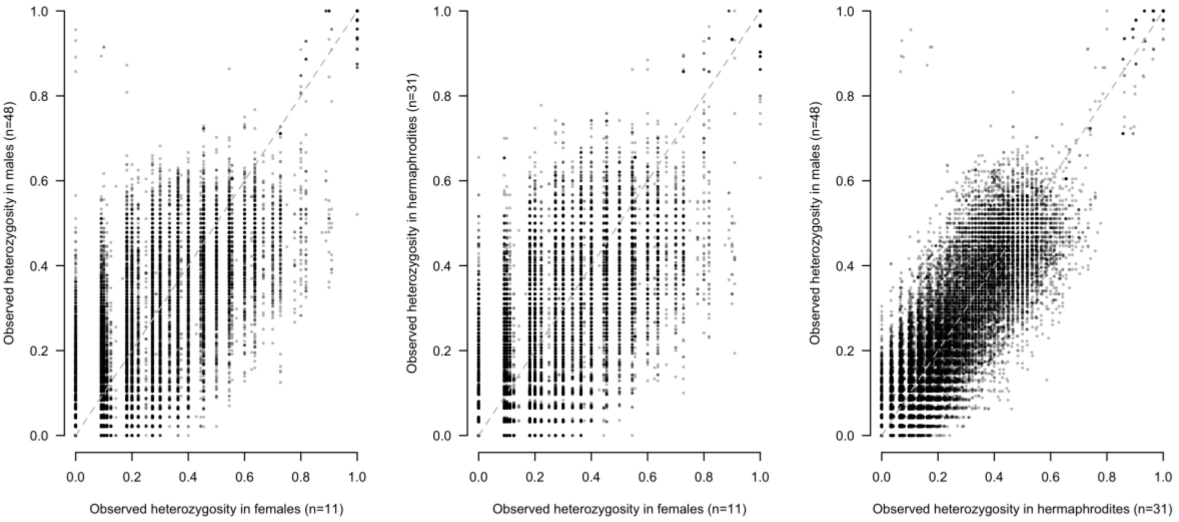


Figure S7 - Locus-specific observed heterozygosity $H_o$ at 20 177 SNPs in *Alviniconcha kojimai* when considering three sex classes: males (*n*=49), females (*n*=11), and morphological hermaphrodites (*n*=31). In the top-left corner of panels A and C, a group of eight SNPs consistently showed strong heterozygosity in males but low heterozygosity in both hermaphrodites and females, and two additional SNPs showed strong heterozygosity in males but low heterozygosity in either hermaphrodites or females.

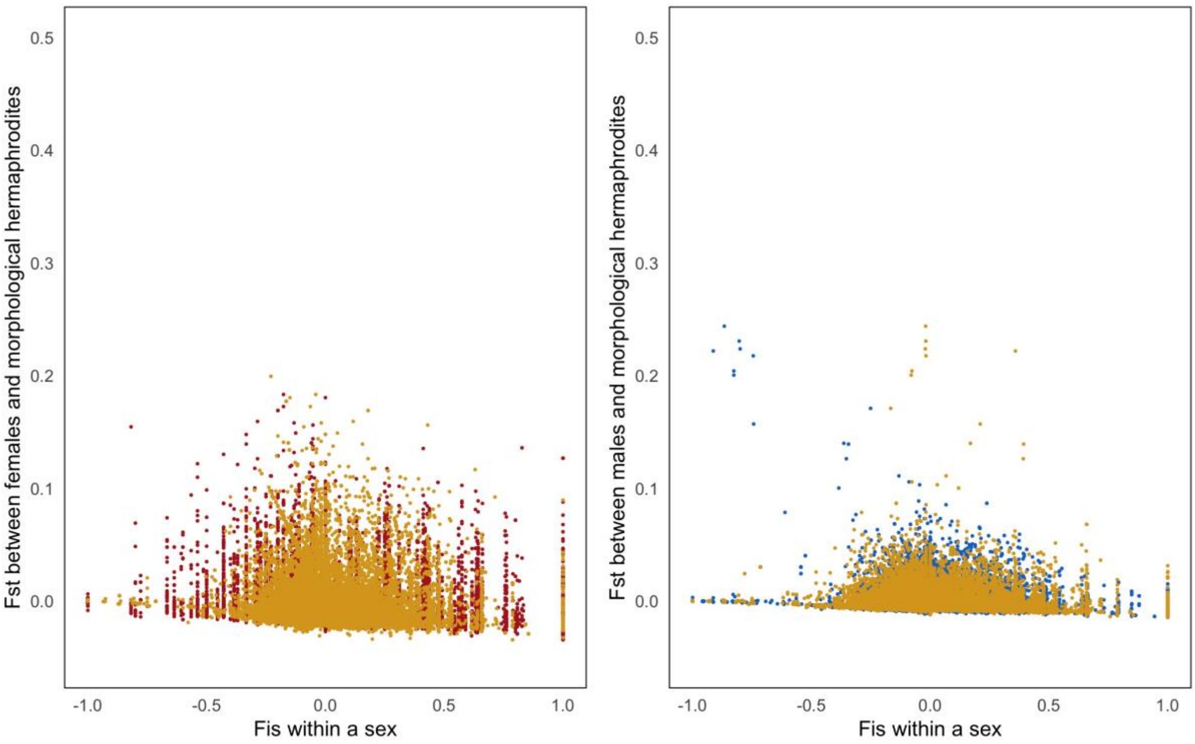


Figure S8 - Locus-specific $F_{ST}$ between males and females and sex-specific $F_{IS}$ for *Alviniconcha kojimai* when considering three sex classes: males (*n*=49, blue), females (*n*=11, pink), and morphological hermaphrodites (*n*=31, orange). Each dot shows observed values for a sex-specific $F_{IS}$ and $F_{ST}$ between two sex classes. The blue dots with strongest $F_{ST}$ between males and morphological hermaphrodites and negative male $F_{IS}$ are potential sex-linked SNPs with contrasted allelic frequencies on X and Y chromosomes in a system where the XX genotype triggers the development of the female or morphological hermaphrodite phenotype while XY triggers the development of the male phenotype (see main text).
